# Recent evolution of large offspring size and post-fertilization nutrient provisioning in swordtails

**DOI:** 10.1101/2023.12.15.571831

**Authors:** Cheyenne Y. Payne, Derek Ly, Rebecca A. Rodriguez-Soto, Daniel L. Powell, Nim D. Robles, Theresa Gunn, John J Bazcenas, Abby J. Bergman, Alexa Pollock, Ben M. Moran, Julie C. Baker, David Reznick, Molly Schumer

## Abstract

Organisms have evolved diverse reproductive strategies that impact the probability that their offspring survive to adulthood. Here, we describe divergence in reproductive strategy between two closely related species of swordtail fish (*Xiphophorus*). Swordtail fish and their relatives have evolved viviparity: they have internal fertilization and give birth to fully developed fry. We find that one species, *X. malinche*, which lives in high-elevation environments, has evolved larger offspring than its closest relative *X. birchmanni* and dwarfs the offspring size of other species in the genus. The larger fry of *X. malinche* are more resilient to starvation than their *X. birchmanni* relatives, hinting that the evolution of large offspring size may be an adaptation to the particularly challenging environments in which *X. malinche* are born. We find evidence that *X. malinche* achieves larger offspring size in part by continuing to provision their offspring over the course of embryonic development after fertilization, the first time this process has been documented in the *Xiphophorus* genus. Moreover, we observe differential regulation in the ovary of genes associated with maternal nutrient provisioning in other species that use this reproductive strategy. Intriguingly, these reproductive differences may drive an asymmetric hybrid incompatibility, since *X. birchmanni* mothers pregnant with F_1_ embryos give birth to premature and stillborn fry at an exceptionally high rate.

## Introduction

Sexually reproducing organisms vary vastly in their investment in their offspring. In some species, investment stops prior to fertilization, whereas in others investment and parental care continue into adulthood (Gross 2005; Klug and Bonsall 2014; Furness and Capellini 2019). In vertebrates, for example, reproductive strategies range from broadcast-spawning millions of eggs (e.g. Atlantic cod; Roney et al. 2018) to a parent raising a single offspring over decades (e.g. orca whales; Weiss et al. 2023). Though the degree of parental investment is generally positively correlated with the probability of offspring survival (Brockelman 1975; Einum and Fleming 2000), there are well-documented tradeoffs between reproductive investment and life history traits like parent survivorship, frequency of reproduction, the number of offspring per reproductive cycle, and the total number over the parent’s lifetime (Smith and Fretwell 1974; Brockelman 1975; Stearns 1989; Einum and Fleming 2000; Jørgensen et al. 2011; Roney et al. 2018).

One measure of reproductive investment that has been studied in diverse organisms across the tree of life is offspring size. Within species, size at birth or weaning is strongly correlated with the probability of offspring survival to reproductive maturity (although there are some exceptions; see Kaplan 1992). The mechanisms through which larger offspring achieve better outcomes are incompletely understood but appear to be diverse and vary between species (such as by avoiding size-dependent mortality, starvation, disease, and conspecific competition; Rollinson and Hutchings 2013; Pettersen et al. 2022). Researchers have speculated that these factors could explain the evolution of differences in offspring size across species.

Another mechanism for increasing offspring survival that has recurrently evolved is viviparity, or internal development of embryos. Viviparity reduces size-dependent mortality of eggs and embryos (Jørgensen et al. 2011) and can offer improved environmental conditions for developing offspring. For some species, it provides an opportunity to directly provision nutrients to developing offspring (Griffith and Wagner 2017). While viviparous species typically provide some level of nutrition to their offspring in the form of yolk in the egg, a strategy known as lecithotrophy, some provision nutrients both before and after fertilization, a strategy known as matrotrophy (Wourms et al. 1988). This post-fertilization provisioning is mediated through physical interfaces between parent and offspring tissue, such as the mammalian placenta (Meredith et al. 2011). While the complex mammalian placenta evolved once over 100 million years ago (Meredith et al. 2011), maternal provisioning structures of varying complexity have evolved over 130 times across vertebrates (Blackburn 2015; Whittington et al. 2022). Lineages in which nutrient provisioning evolved recently may be especially useful in understanding the pressures that drive post-fertilization nutrient provisioning.

Poeciliid fishes offer an opportunity to study the evolutionary drivers of viviparity, nutrient provisioning strategies, and variation in offspring size. The common ancestor of poeciliids evolved internal fertilization and live birth of juvenile fish, or fry (Pollux et al. 2009), and species vary widely in the degree of post-fertilization nutrient provisioning (Reznick et al. 2002; Pires et al. 2010). Counterintuitively, the evolution of offspring size and matrotrophy are often decoupled in poeciliids. While it might be expected that provisioning more nutrients during development would lead to larger offspring at birth, empirical data have yielded no consistent association across taxa (Olivera-Tlahuel et al. 2015; Meiri et al. 2020; Furness et al. 2021).

However, potential links between offspring size and matrotrophy have been difficult to disentangle because offspring size is often strongly impacted by maternal traits like age, size, and condition (Berkeley et al. 2004; Marshall and Keough 2007; Hagmayer et al. 2018; but see Marshall et al. 2010). Ecological factors are also important drivers of offspring size, independent of post-fertilization nutrient provisioning (Reznick et al. 1996*a*; Jennions et al. 2006; Riesch et al. 2010; Pollux and Reznick 2011; Schrader and Travis 2012; Leips et al. 2013). For example, offspring of lecithotrophic *Poecilia* from environments with low predation and high competition are 50% larger at birth than those from high predation, low competition environments (Reznick 1982*a*; Reznick and Endler 1982; Reznick and Bryga 1987; Reznick et al. 1996*c*; Bashey 2006*a*, 2008; Jørgensen et al. 2011), presumably as a result of mothers investing in larger eggs. Similar environmental conditions appear to promote larger offspring size in some matrotrophic species (Schrader and Travis 2012; Leips et al. 2013; but see Reznick et al. 1996*b*) but not in others (Pollux and Reznick 2011).

Here, we address questions about the evolution of offspring size and matrotrophy in a group of livebearing fish that are an important evolutionary and behavioral model system. *Xiphophorus* species inhabit dramatically different ecological environments, from valley rivers nearly at sea level to mountain streams in the Sierra Madre Oriental in central México. Past work on a handful of species has suggested that although all *Xiphophorus* species are livebearing, they do not invest in their offspring post-fertilization (i.e. they are lecithotrophic; Constantz et al. 1989). However, this hypothesis has not been evaluated on a genus-wide scale, and data is particularly lacking in species living in extreme environments for the group (Morris and Ryan 1992; Pollux et al. 2014). There is no data available on variation in offspring size within or between species.

We find that *X. malinche* have evolved exceptionally large offspring and explore the mechanisms and pressures potentially driving the evolution of this trait. By characterizing the change in embryo size over development in the lab and wild for the closely related northern swordtail species *X. malinche* and *X. birchmanni*, we find evidence that *X. malinche* achieves larger offspring size in part through post-fertilization maternal nutrient provisioning. Using a combination of approaches, including evaluating expression at genes that have been repeatedly co-opted for post-fertilization maternal nutrient provisioning (Guernsey et al. 2020), we investigate some of the potential mechanisms underlying this reproductive strategy in *X. malinche*. Notably, *X. malinche* and *X. birchmanni* naturally hybridize (Culumber et al. 2011), which we leverage to study the link between genome-wide ancestry and offspring size. Finally, we explore the possibility that a conflict between maternal nutrient provisioning and offspring nutrient demands over development underlies asymmetric hybrid inviability between these two species. This observation is particularly exciting given ongoing hybridization between these two species in nature.

## Methods

### Measuring newborn fry size across Xiphophorus species

To compare the size of newborn fry in several *Xiphophorus* species, broods were collected in the laboratory on the day they were born. All tanks with gravid females were inspected multiple times a day for fry. Upon collection, all fry from a brood were photographed in shallow water (∼2 cm deep) from overhead for a dorsal view of each fry. Photos were imported to ImageJ2 and two measurements, standard length and head width, were made using the line measurement tool (Supp. Fig. 1). We measured both traits for fry from five *Xiphophorus* species – *X. birchmanni, X. malinche, X. cortezi, X. pygmaeus*, and *X. variatus*, and from *X. malinche x X. birchmanni* F_1_, F_2_, and lab-reared natural hybrids descended from individuals collected from the Calnali Low wild population. Note that we report newborn fry size for the *X. malinche x X. birchmanni* F_1_ cross direction only, as F_1_s from the reciprocal *X. birchmanni x X. malinche* are rarely carried to term (see below). Given that fry were collected from tanks with multiple gravid females (*X. birchmanni* and *X. malinche* experience high levels of stress when singly housed), we were not able to collect covariates such as mother standard length. However, because average mother size is similar between species (on average, 4.04 cm for *X. malinche* and 3.94 for *X. birchmanni* reared in common conditions; see Supplementary Information 1), we expect that our large sample sizes (*X. malinche* broods N=35 and *X. birchmanni* broods N=48) will not be dominated by effects of individual mother size. Since broods varied in size, we averaged the standard length and head width for each brood and compared means with a Wilcoxon rank sum exact test. See Table S1 for raw fry standard length and head width and Table S2 for statistics. Using lab-reared *X. birchmanni*. *X. malinche,* and hybrid fry, we also obtained an estimate of broad-sense heritability (H^2^) of offspring standard length at birth (see Supplementary Information 2 for methods).

### Measuring embryo weight across developmental stages

To compare embryo size throughout development in each species, we measured the dry weights of embryos across developmental stages from wild-caught females, collected with baited minnow traps. We collected a total of 38 pregnant and 11 nonpregnant females (with fully-yolked eggs) from the *X. malinche* source population Chicayotla on the Río Xontla (1,003 m elevation; 20°55′27.24″N 98°34′34.50″W) across two seasons (45 in May 2022 and 4 in August 2020), 59 pregnant and 51 nonpregnant females from the *X. birchmanni* source population Coacuilco on the Río Coacuilco (320 m elevation; 21°5’50.85 N, 98°35’19.46 W) across four seasons (31 in October 2023, 22 in February 2023, 30 in September 2022, and 27 in August 2020), and 19 pregnant females from the *X. cortezi* source population Puente de Huichihuayán on the Río Huichihuayán (89 m elevation; 21°26’9.95 N 98°56’0.00 W) in one season (February 2023). In practice, the sample of 19 females from *X. cortezi* was insufficient to capture developmental profiles because we had ≤1 brood for many stages, so we limit our discussion of these results to the supplement (Supplementary Information 3). Each mother was euthanized with MS-222, cut from the anal pore to the gills so as to expose the brood for fixation, and stored in 95% ethanol. Consistent with previous reports for *Xiphophorus* (Kindsvater et al. 2012; Furness et al. 2019), we found that all embryos in a brood were at roughly the same developmental stage.

Embryos were staged following the numerical scoring guidelines described by Reznick 1981. Briefly, stages range from 0-50 (in intervals of 5), where stage 0 describes yolking or fully yolked unfertilized eggs and stage 50 describes fully developed fry with a closed pericardial cavity (see Table S3). Importantly, these stages are categorical based on key morphological features of developing embryos and do not reflect constant time intervals throughout development, as the temporal interval varies. It is typically difficult to discern whether stage 0 eggs are fully yolked or are still yolking. Therefore, we noted whether unfertilized eggs had an even distribution of lipid droplets on the surface of the egg and were evenly sized within their brood, which are the best visual indicators of completed yolking.

Embryos and eggs were separated from the ovarian follicle, staged under a microscope, and then dried following methods described in D. Reznick (1981). The ovarian tissue and each staged embryo and egg were placed in individual microcentrifuge tubes that were dried in a laboratory oven overnight at 65°C, and then dry weighed using a standard scale sensitive to 10^-4^ grams (Table S4).

### Comparison of embryo weight over development between species

Maternal size has previously been shown to strongly correlate with offspring size in poeciliids, including in *X. birchmanni* (Jørgensen et al. 2011; Kindsvater et al. 2012; Hagmayer et al. 2018), as have brood size and environmental variables like collection season (Kindsvater et al. 2012). In our wild collections, pregnant mothers ranged in length from 3.4-5.7cm for *X. malinche* and 2.8-5.2cm for *X. birchmanni*, and brood size ranged from 3-36 and 4-44 embryos, respectively. Therefore, to evaluate which variables impact embryo dry weight other than our variables of interest (species, stage, and the interaction between species and stage), we used R stats::step to compare models that included mother standard length, brood size, and season in an AIC framework. Collection dates were binned into two seasonal categories describing water temperature (Supp. Fig. S11): the “warm” season includes April-October months and the “cold” season includes November-March months. Mother standard length and season were selected and included as covariates with fixed effects. We confirmed that there was not a significant interaction between species and mother standard length before proceeding with analysis. We also note that, reassuringly, the overall patterns in our results are similar with and without standard length and season covariates included in the model. For the full analysis, we also included brood ID as a random effect, which is unique to each female and is species-specific. Therefore, we fit the following mixed linear model with R lme4::lmer: embryo dry weight (g) ∼ species + stage + species:stage + season + mother standard length + (1 | brood ID). Pairwise comparisons between mean embryo dry weight for each stage were made with an ANOVA and Tukey post-hoc test with R emmeans::emmeans (Table S6).

We visualized differences in embryo size after accounting for covariates by plotting the partial residuals of embryo dry weight for each stage with R visreg::visreg (Breheny and Burchett 2017), split by species. We also plotted the raw data (i.e. without accounting for covariates) and reassuringly, the overall patterns were similar (Fig. S5).

As noted above, because it is difficult to determine whether stage 0 eggs are fully-yolked, we omitted these eggs from the developmental profile plots, but separately plot the stage 0 eggs that appeared to have even lipid distribution and size within their brood (Table S5). We used the subset of unfertilized eggs that met these criteria to compare pre-fertilized egg weight between species but note that the distribution of these dry weights likely skews lower than the true pre-fertilized egg weights for both species (see Table S5). We fit a mixed linear model of egg dry weight by species with mother standard length and season as selected covariates and brood ID as a random effect, performed pairwise comparisons with an ANOVA and Tukey post-hoc, and plotted partial residuals of egg dry weights as above.

### Assessing the relationship between genome-wide ancestry and embryo and ovary size

We were curious whether genome-wide ancestry proportion in natural hybrid mothers might predict embryo size and ovary weight, given that this trait is heritable in lab-raised hybrids (see Results). We sampled 34 pregnant mothers from a natural hybrid population on the Río Calnali on a single collection date (the Calnali Low population, elevation 905 m). *X. malinche* ancestry proportions ranged from 0.28 to 0.71 in these individuals (see Table S7 and Supplementary Information 4). Embryos and ovary tissue were dissected, staged, and dry-weighed as described in the previous section. As above, we used R stats::step to choose an appropriate model for testing the relationship between embryo dry weight, as well as ovary dry weight, and mother ancestry proportion. The selected mixed linear model included brood size, stage, and the interaction between stage and ancestry proportion, with brood ID as a random effect. We evaluated significance with a likelihood ratio test using R stats::anova to compare the full model and a reduced model without ancestry proportion and its interaction with stage. We also evaluated whether mother mitochondrial genotype predicted embryo or ovary dry weight (see Supplementary Information 4). We calculated and plotted partial residuals of embryo dry weight after accounting for covariates (e.g. ancestry proportion) with R visreg::visreg.

### Calculating Matrotrophy Index

The degree of post-fertilization nutrient provisioning varies along a continuum, but researchers have described two general provisioning strategies: lecithotrophy and matrotrophy (Pollux et al. 2009). A simple quantitative metric for distinguishing these strategies and measuring the level of post-fertilization provisioning is the Matrotrophy Index, which is the dry weight of the fully-developed embryo divided by the dry weight of the fully-yolked egg. A Matrotrophy Index < 1 indicates that the embryo has decreased in size over development (lecithotrophy) while a value ≥ 1 indicates that the embryo has maintained or increased in size, indicative of some level of maternal provisioning over development (matrotrophy). Reported Matrotrophy Indices within poeciliids range from 0.45 to 103 (Furness et al. 2021). Note that a large Matrotrophy Index does not necessarily correspond to large offspring size at birth.

We calculated Matrotrophy Index for *X. malinche* and *X. birchmanni* as the dry weight of a fully-developed embryo divided by the dry weight of an early-stage embryo, averaging dry weights for embryos of the same stage within a brood. Since it is difficult to identify fully-yolked eggs, we calculated Matrotrophy Index with dry weight measurements from stage 10 embryos. This analysis choice is conservative in the context of our study, since it will give an underestimate of Matrotrophy Index in matrotrophic species. Since multiple variables impact embryo dry weight, we calculated Matrotrophy Index using both the raw averages (Table S4) and the partial residuals of average embryo dry weights (Table S8) after accounting for season and mother standard length as covariates as above (with R stats::lm). We describe the impact of different calculations with raw and partial residual dry weights on our estimate of Matrotrophy Index in Supplementary Information 3.

### Multifactorial artificial insemination crosses

To compare results from wild-caught females to females raised in controlled lab conditions, we artificially inseminated lab-raised *X. malinche* and *X. birchmanni* mothers following the *Xiphophorus* Genetic Stock Center Protocol to create the following crosses: *X. malinche ♀* × *X. malinche ♂, X. birchmanni ♀* × *X.birchmanni ♂, X. malinche ♀* × *X. birchmanni ♂*, and *X. birchmanni ♀* × *X. malinche ♂*. Artificially inseminated females from each cross (N=10-15) were reared in 200-gallon outdoor tanks at Stanford University until females were visually determined to be late in pregnancy in April-May 2023, at which point all fish were sedated with MS-222 and euthanized by severing the spinal cord. Embryos were dissected, and each embryo was assigned a developmental stage following Table S3. Dry weight of each embryo and the ovarian tissue were measured as described above (see Table S9). Because we were interested in evaluating cross-level differences in late-stage embryo size, we subsampled stage 30-50 embryos (from 2 broods for all crosses except *X. birchmanni ♀* × *X. malinche ♂*, for which there was only 1 brood; see Table S9). We found that for this dataset, due to the small sample size, mother standard length and brood ID were perfectly correlated, and including brood ID as a random effect caused singularity in the model. Therefore, we fit a simple linear model including species, stage, brood size, and mother standard length as selected covariates with R stats::lm, ANOVA, and Tukey post-hoc test with R emmeans::emmeans (see Table S10), and plotted partial residuals of dry weights with R visreg::visreg. Reassuringly, the partial residual results are the same whether brood ID is included as a random effect in the model or not. We also made crosses between individuals of the same species from different populations for comparison (Table S11-12 and Supplementary Information 5).

To look for evidence of anatomical differences between species at the maternal-offspring interface that could be associated with nutrient provisioning (i.e. thicker ovarian follicle and interacting embryo and maternal tissue), we prepared histological slides of ovaries from both nonpregnant and late-stage (stage 35-40) pregnancy females from the pure *X. malinche* and *X*. *birchmanni* crosses, as well as both F_1_ crosses. Whole ovaries containing eggs or embryos were carefully dissected and fixed in 10% formalin. Ovaries were paraffin-embedded, sectioned, and stained with Hematoxylin & Eosin by the Histo-Tec Laboratory (Hayward, CA). We digitally scanned the stained slides through the Human Pathology Histology Services Laboratory at Stanford University for morphological analysis of the ovary sections. See Supplementary Information 6 for methods quantifying morphological differences. Additionally, we verified the genetic origin of ovary follicle tissue in *X. malinche* mothers carrying F_1_ offspring (Supplementary Information 7).

### Differential gene expression and co-expression network analysis

Previous work in livebearers has identified several key genes that are differentially expressed between the ovarian tissue of pregnant lecithotrophic and matrotrophic species (Jue et al. 2018; Guernsey et al. 2020). To broadly compare gene expression between *X. malinche*, *X. birchmanni,* and *X. cortezi,* we sequenced ovarian tissue from mothers from two groups: those with fully-yolked eggs and those with embryos in mid-to late-development.

Because we were interested in species-level rather than environmentally-dependent differences, we collected both wild and lab-reared females for this analysis. For wild-caught fish, we collected two *X. malinche* and two *X. birchmanni* pregnant females with stage 40-50 embryos from the Chicayotla and Coacuilco localities, respectively. Additionally, we sampled at least three “early” pregnancy females (unfertilized fully-yolked or stage ≤ 5) and at least three “late” pregnancy females (stage 25-45) from our lab populations. For both sets of samples, fish were euthanized by rapidly severing the spinal cord with a scalpel and dissecting the mother’s body cavity from anal fin to gills. Wild-caught fish were stored in RNAlater and ovaries were dissected later. For lab collected fish, whole ovaries were dissected immediately following euthanasia and preserved in RNAlater. See Table S13 for the full list of samples with collection dates and developmental stages of embryos.

Ovarian tissue and embryos were carefully separated. See Supplementary Information 8 and Table S14 for differential gene expression analysis of embryos. RNA was extracted from ovarian tissue from each mother using the Qiagen Mini RNAeasy kit. RNAseq libraries were prepared with unique barcodes for each sample using a KAPA mRNA HyperPrep Kit, pooled, and sequenced. The wild-caught samples and half of the lab samples were sequenced on one Illumina HiSeq4000 lane and the rest of the lab samples were sequenced on one Illumina NovaSeq6000 lane for an average of 28 million 150 bp paired-end reads per sample (see Table S13). Raw reads are available under NCBI BioProject PRJNAXXXX. For each sequencing batch, samples were paired to balance species and developmental stages so that batch effects could be statistically accounted for in analysis.

We followed the gene expression analysis methods described in Payne et al (2022). We used cutadapt (Martin 2011) and Trim Galore! (Krueger et al. 2021) to trim Illumina adapter sequences and low-quality bases (Phred score < 30) from reads. Using the tool *kallisto* (Bray et al. 2016), we pseudoaligned RNAseq reads to an *X. birchmanni* “pseudoreference” transcriptome generated from the southern platyfish *X. maculatus* genome (Schartl et al. 2013). The *X. maculatus* genome assembly is well-annotated and mapped to Ensembl gene IDs with associated Gene Ontology terms (Wittbrodt et al. 1989; Schartl et al. 2013). Raw transcript counts were converted to gene-level counts to evaluate gene expression; note that the *X. birchmanni* transcriptome contains only a single transcript per gene.

For differential gene expression analysis, we combined all lab- and wild-collected ovarian follicle samples into a single dataset, using a design formula that included species (*X. malinche*, *X. birchmanni*, or *X. cortezi*) and pregnancy category (early or late) as a grouped interaction, origin (lab or wild), and library preparation/sequencing batch. Briefly, using R DESeq2::DESeq (Love et al. 2014), we normalized gene counts by library size, estimated within-experimental group dispersion, fit a negative binomial generalized linear model, and tested significance with a Wald test. Shrunken log-fold changes were calculated with the ashr::ashr shrinkage estimator (Stephens 2017). Extreme outlier and low count genes were removed. Genes were considered significantly differentially expressed between species at an FDR-adjusted p-value<0.05. Out of a total of 19,176 genes in the *X. birchmanni* pseudoreference transcriptome, 96% are expressed in at least one species. Expression results from this analysis can be found in Table S15.

To explore biological pathways enriched in genes that were differentially expressed between species in ovarian tissue, we performed Gene Ontology enrichment analysis. We used R biomaRt (Durinck et al. 2009) and GOstats (Falcon and Gentleman 2007) to match *X. maculatus* Ensembl gene IDs with GO terms. We created a “universe” of all genes analyzed with DESeq2, which was used as the reference set of genes for testing category enrichment. Using R GSEABase::hyperGTest (Morgan, Martin et al. n.d.), we ran a hypergeometric test to identify overrepresented GO terms (p-value<0.05) in the set of genes that were significantly differentially expressed between species and developmental stage (Table S16). We also used the R tool WGCNA (Langfelder and Horvath 2008) to cluster co-expressed genes and identify clusters highly correlated with species and developmental stage as a grouped variable. Detailed WGCNA methods are in Supplementary Information 9 and full results are in Tables S17-18.

### Immunostaining of prolactin expression in ovarian tissue

Ovaries were fixed, processed, and sectioned as described above. Briefly, sections were deparaffinized and dehydrated through xylenes and a graded ethanol series. Slides were blocked for 1h (at room temperature, hereafter RT) in gelatin block, then incubated with anti-prolactin primary antibody (rabbit, Abcam EPR19386, 1:200) overnight (4C). Slides were washed, peroxide blocked for 30m (RT) then incubated with biotinylated goat anti-rabbit secondary antibody (Jackson ImmunoResearch 111-065-144, 1:5000) for 1h (RT). Slides were washed, then incubated for 30m with VectaShield Elite ABC reagent (Vector Labs PK-6100; RT). Prolactin signal was amplified with TSA-Cy3 (Akoya Biosciences NEL744001KT) for 6m (RT) and counterstained with DAPI (Thermo Scientific 62248, 1:2000) and mounted with Prolong Gold Antifade (Thermo Scientific P36930). See complete methods in Supplementary Information 10.

Images were acquired using NIS Elements software v4.30.02 on a Nikon Ti Eclipse inverted microscope equipped with an ASI MS-2000 motorized linear XY stage, Yokogawa CSU-W1 single disk (50mM pinhole) spinning disk unit, Andor Zyla 4.2 (6.5mM pixel size) scMOS camera, and 10x/0.45 NA or 20X/0.75 NA Nikon PlanApo Lambda air objectives (see Supplementary Information 10 for complete imaging specifications). Data was saved in .nd2 format and manually processed in ImageJ.

### Exploring ecological differences in X. birchmanni and X. malinche habitats

One striking environmental difference between *X. birchmanni* and *X. malinche* habitats is water temperature, driven by elevation differences (Fig. S10). In addition, many ecological factors such as primary productivity covary with temperature. While we investigated direct responses to temperature in *X. birchmanni* and *X. malinche* fry (Supplementary Information 11 and Table S19), we were also interested in responses to the indirect effects of temperature, such as food availability.

Given that *X. malinche* populations likely experience lower food and nutrient availability, we predicted that newborn fry may be more likely to face starvation conditions than *X. birchmanni*, and that large offspring size may have evolved as an adaptation to these conditions. To directly test this, we compared starvation tolerance between newborn *X. malinche* and *X. birchmanni* fry in the lab. We collected four broods of fry from each species and split individuals in a brood evenly between one experimental and one control tank. Fry in the control tanks were fed brine shrimp twice daily for three days while fry in experimental tanks were not fed for those three days. Observations of physical state and behavior were recorded twice per day at 10:00 and 17:00 during the laboratory light period (09:00-18:00). At the end of the three-day trial, fry were euthanized with MS-222, photographed so that standard length could be measured with ImageJ, and prepared for total lipid extraction to measure differences in dry weight and fat content between control and experimental fry (see below). After model selection, we fit a mixed linear model (R nlme::lme) for standard length, including species, treatment, and their interaction as fixed effects and brood ID as a random effect. Pairwise comparisons between the interaction of species and treatment were made with an ANOVA and Tukey post-hoc test with R emmeans::emmeans and partial residuals were calculated and plotted with lme4::lmer and visreg::visreg. See Table S20 for raw data and Table S21 for statistics.

### Lipid extraction and fat content analysis of wild-caught females and lab-born fry

*Xiphophorus* species were thought to reproduce year-round. However, in sampling natural *X. malinche* and *X. birchmanni* populations for this project we detected clear evidence for seasonality in breeding in *X. malinche* populations (see Results). To more systematically evaluate this, we opportunistically collected pregnancy rates for collections made in August 2020, May 2022, September 2022, and February 2023 from the Chicayotla *X. malinche* and Coacuilco *X. birchmanni* populations (Table S22).

Dry weight and body fat content are indirect measures of nutrient availability in a fish’s environment. We dry weighed and extracted total lipids from wild-caught nonpregnant Chicayotla *X. malinche* and Coacuilco *X. birchmanni* adult females caught in September 2022 and February 2023. We also collected dry weight and total lipid data from the fry subject to the food deprivation trials described above. For adult fish, abdominal organs (i.e. digestive, excretory, and reproductive tissue from the abdomen) were dissected out to avoid extracting lipids from undigested food. Fry were left intact. Fish were placed on a tray wrapped with foil and dried in a drying oven set to 65℃ for five days. After desiccation, dried fish were weighed to obtain dry weight prior to lipid extraction (weight_1_). To extract lipids, each dried fish was placed in a 5mL glass vial of petroleum ether. After 24 hours, the petroleum ether was drained and replaced with fresh petroleum ether. This process was repeated once more to ensure complete lipid extraction, for a total of three petroleum ether washes. After the third wash, fish were dried at 65℃ for 24 hours and then dry weighed to obtain dry weight after lipid extraction (weight_2_). Fat content percentage was calculated as:

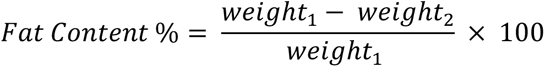

Differences in fat content for nonpregnant females for each season were evaluated with a Student’s two-sided t-test. Differences in dry weight and fat content between species and treatments for food deprivation trial fry were evaluated with the same methods described for fry standard length (see above), with the same model selected (Tables S20-21).

### Measuring viability by cross direction

In addition to comparing embryo size in F_1_ hybrids, we quantified differences in survival of F_1_ fry as a function of cross direction. We used additional F_1_ crosses generated in the lab to do so. For these crosses, we monitored females twice daily, and to reduce the risk of fry predation, we provided low-density and high-cover conditions for females that were morphologically identified as being close to giving birth. New broods were morphologically scored for being full-term or premature, and stillborn fry were identified and counted. Tanks of collected fry were monitored twice daily. For crosses with high mortality rates, survival to two weeks was recorded for surviving fry (Table S23).

## Results

### X. malinche newborn fry are exceptionally large

To evaluate variation in offspring size at birth across *Xiphophorus*, we measured the standard lengths and head widths of newborn fry from five species: *X. birchmanni, X. malinche, X. cortezi, X. pygmaeus*, and *X. variatus* (Fig. 1A, Supp. Fig. S1-2, Table S1). We found significant differences in size across species (Table S2). Notably, *X. malinche* have the largest fry (10.8 mm ± 1.1 mm; adjusted p-value<0.005 for all five comparisons, see Table S2). Despite this difference in offspring size, *X. birchmanni* and *X. malinche* did not consistently differ in the number of offspring per brood (although offspring size varied as a function of mother size and collection season; Supp. Fig. S3). Moreover, adult females of the two species grown in common garden conditions were similar in size (Supp. Fig. S4). Although *X. birchmanni* fry are significantly smaller than *X. malinche* fry, they are larger than fry of other *Xiphophorus* species (Fig. 1; Table S2).

**Figure 1.**
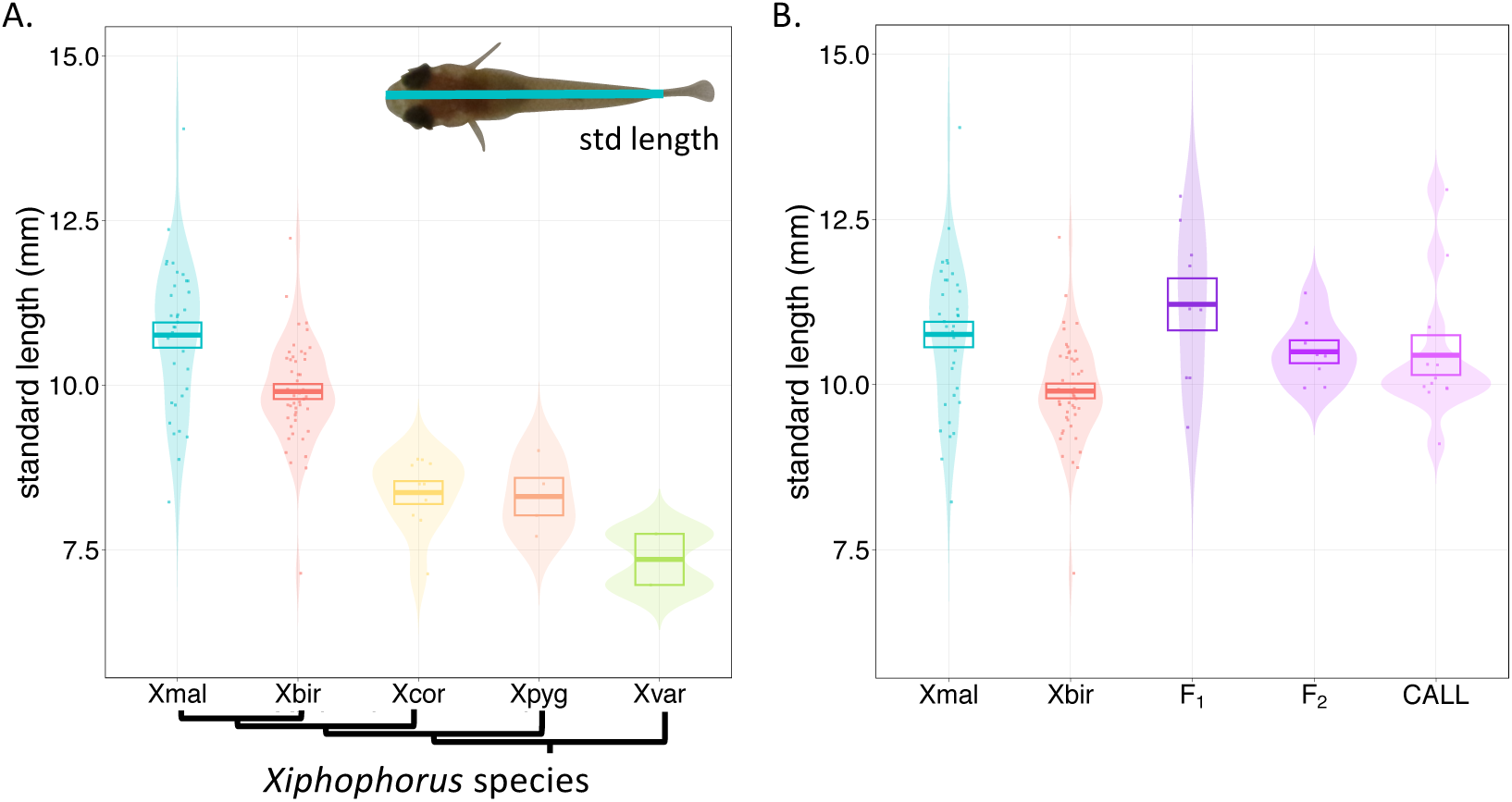
Comparisons of offspring size at birth across *Xiphophorus* species and their hybrids. **A)** Violin plots of standard length (mm) of newborn fry from five *Xiphophorus* species – *X. malinche*, *X. birchmanni*, *X. cortezi*, *X. pygmaeus*, and *X. variatus*. Phylogenetic relationships between species are shown schematically on the x-axis (following (Preising et al. 2022). **B)** Violin plots of standard length (mm) of newborn fry of different groups, including *X. malinche*, *X. birchmanni*, F_1_ hybrids, F_2_ hybrids, and natural hybrids (“CALL” – Calnali Low hybrid population). For both **A** & **B**, each data point represents the average standard length of newborn fry from one brood. For each group, boxes show the mean ± 1 standard error.

We also compared the size of newborn *X. malinche x X. birchmanni* F_1_ and F_2_ fry. We found that F_1_ standard length and head width was *X. malinche*-like and F_2_ and natural hybrid fry were intermediate in size to their parent species (Fig. 1B, Supp. Fig. 2B, Tables S1-2). After accounting for the effects of season and brood ID, we estimated the broad-sense heritability of standard length at birth attributable to ancestry to be 0.25 (see Supplementary Information 1).

### X. malinche mothers provision developing embryos

Given that *X. malinche* fry are significantly larger at birth than *X. birchmanni*, we hypothesized that *X. malinche* embryos are also larger throughout embryonic development. We measured the dry weight of embryos dissected from wild-caught *X. malinche* and *X. birchmanni* pregnant females across developmental stages. First, we estimated the Matrotrophy Index for each species to obtain a quantitative metric of nutrient provisioning during development using stage 10 and stage 50 embryos (Fig. 2A; see Supplementary Information 3). On average by brood, stage 10 embryos weighed 0.0038g for *X. birchmanni* and 0.0040g for *X. malinche*, while stage 50 embryos weighed 0.0025g and 0.0038g, respectively. We note, however, that in practice, embryo weight is impacted by season and mother standard length (see Methods). After accounting for season and mother standard length, we calculated Matrotrophy Indices of 0.66 for *X. birchmanni* and 0.98 for *X. malinche*, respectively. We also attempted to calculate Matrotrophy Index for the sister species to the *X. malinche-X. birchmanni* clade, *X. cortezi*, but our small sample size led to uncertainty in these estimates (Supplementary Information 3).

**Figure 2.**
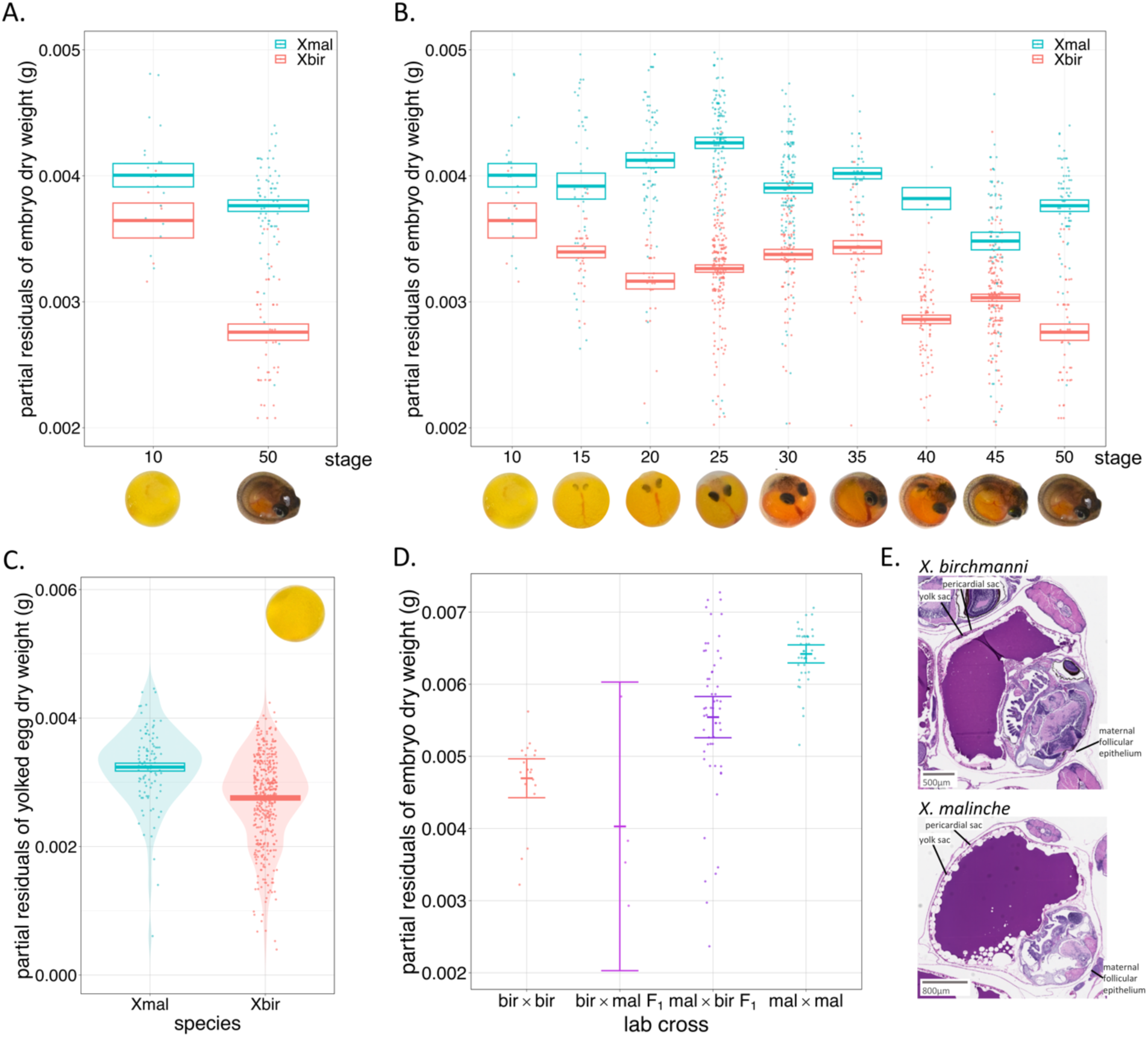
Comparisons of embryo weight throughout development and ovarian structures between *X. birchmanni* and *X. malinche*. **A)** Dry weights of wild-caught *X. birchmanni* and *X. malinche* embryos do not significantly differ shortly after fertilization but are substantially different in weight just before birth (as well as at birth, see Fig. 1). Plot shows partial residuals of stage 10 and stage 50 embryo dry weights correcting for season and mother standard length as covariates and brood ID as a random effect. Points show the partial residuals and boxes show mean ±1 standard error, colored by species (*X. birchmanni/*Xbir – pink and *X. malinche/*Xmal – blue). The y-axis was limited to 0.002-0.005g to show the mean differences between species by stage more clearly; however, several points were cut out of the plot frame. The full plot including all partial residual points is shown in Fig. S5A. **B**) Developmental size profiles for wild-caught *X. birchmanni* (Xbir – pink) and *X. malinche* (Xmal – blue) show that the difference in embryo size between these species may fluctuate throughout development (or be impacted by variation in sampling), but appears to diverge after stage 15. Developmental stage is plotted on the x-axis (with images of *X. malinche* embryos at corresponding stages) and the partial residuals of average embryo dry weight per brood (after correcting for maternal size and collection season) is plotted on the y-axis. Points show the partial residuals and boxes show mean ±1 standard error, colored by species. **C**) Though difficult to accurately stage (see Methods), unfertilized, fully-yolked eggs do not significantly differ in size between species (p-value=0.18). Violin plot compares stage 0 (fully-yolked but unfertilized) eggs between wild-caught *X. birchmanni* and *X. malinche*. Points show the partial residuals and boxes show mean ±1 standard error, colored by species. **D**) Patterns observed in **B** are replicated in stage ≥30 lab-crossed parental species raised under common conditions, with *X. birchmanni* embryos being significantly smaller than *X. malinche* later in development (p-value=0.0005). Given the small sample size of late-stage F_1_ hybrids from *X. birchmanni* mothers (bir×mal F_1_), we have limited power to evaluate whether they differ in size from the other crosses. F_1_ hybrids from *X. malinche* mothers (mal×bir F_1_) trend towards being smaller than pure *X. malinche* fry (p-value=0.08), hinting that the trait could be impacted by both genetic and maternal effects. **E)** Histological slides stained with Hematoxylin & Eosin of *X. birchmanni* (left) and *X. malinche* (right) ovaries derived from females pregnant with late-stage embryos (stage 35 and 40, respectively) are structurally similar.

Consistent with their Matrotrophy Index estimate, we found that *X. birchmanni* embryos show a lecithotrophic developmental profile, where embryos lose weight overall across development (Fig. 2B). Notably, *X. birchmanni* embryonic size appears to change nonlinearly through development. By contrast, *X. malinche* embryos maintain roughly the same dry weight throughout development. These species do not differ in embryo dry weight early in development (p-value=0.38 at stage 10; Fig. 2A) nor at the time of fertilization (p-value=0.18, Fig. 2C; but note methodological issues with this stage discussed above). However, they significantly diverge in embryo dry weight at stage 20 (p-value=0.008), and ultimately differ substantially by late development (p-value=0.002 at stage 50; Fig. 2A-B; see Table S5 for statistical comparisons by stage). Therefore, the maintenance of weight over development by *X. malinche* embryos suggests that there is some level of post-fertilization nutrient provisioning from *X. malinche* mothers.

### Late-stage F_1_ embryos in the X. malinche x X. birchmanni cross are intermediate in size

To compare the size of late-stage embryos (stage ≥30) from the reciprocal F_1_ crosses to pure parental crosses, we made controlled crosses between *X. malinche* and *X. birchmanni* (see *Methods*). Consistent with our measures from wild-caught broods, lab-bred *X. malinche* embryos are significantly larger than *X. birchmanni* embryos (p-value<0.0005). We find that F_1_ embryos from the *X. malinche* mother and *X. birchmanni* father cross direction trend towards being smaller than *X. malinche* embryos (p-value=0.08) but span the size range of both within-species crosses. This hints that in a *X. malinche* maternal environment, both maternal effects and offspring genotype could contribute to offspring size at birth. For F_1_ embryos with *X. birchmanni* mothers, we were unable to sample sufficient numbers of embryos to confidently compare size across groups (Fig. 3C; see Table S10 for summary statistics).

**Figure 3.**
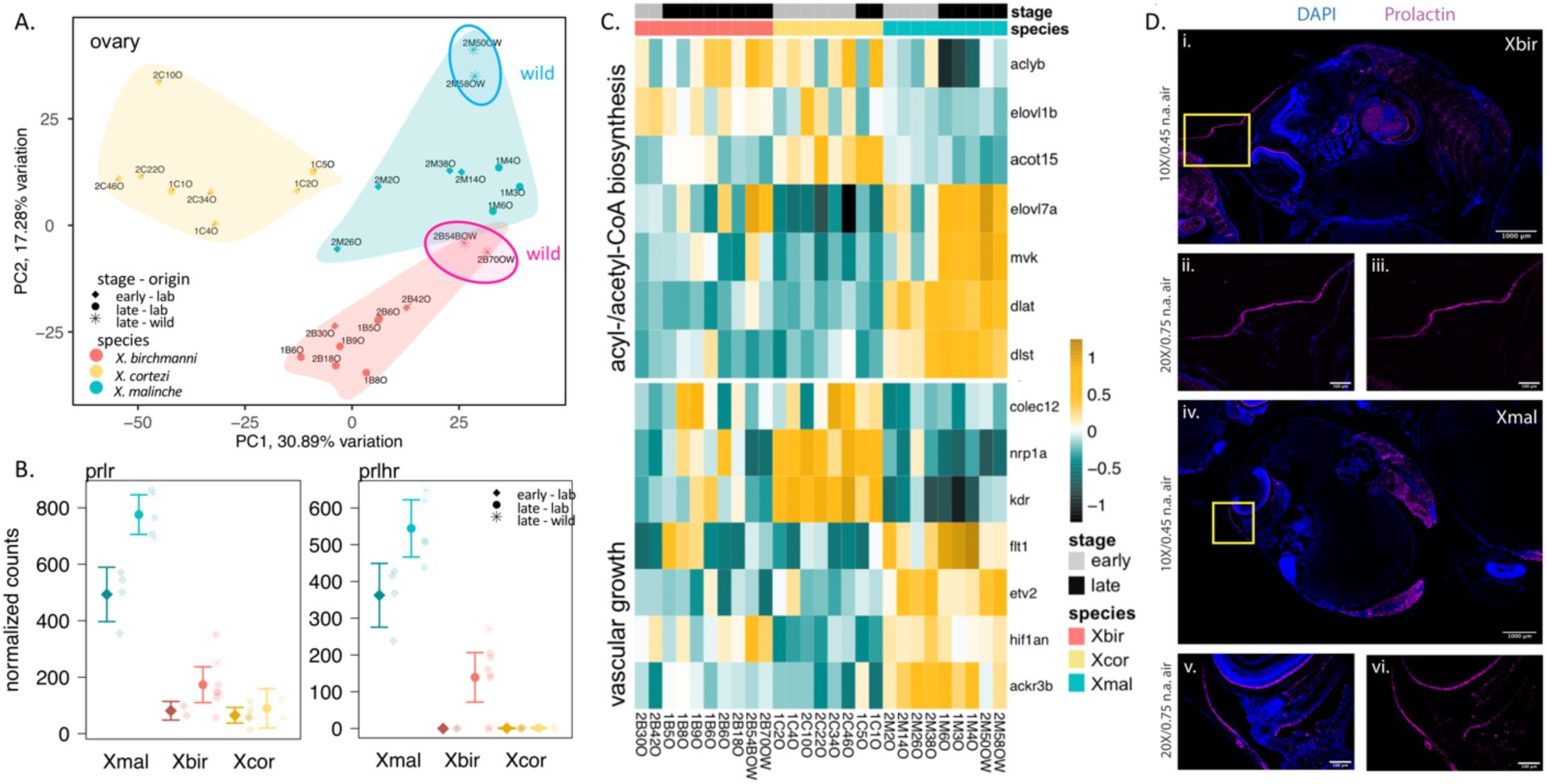
Gene expression analysis hints at a role for prolactin, fatty acid biosynthesis, and vascular growth pathways in nutrient provisioning post-fertilization in *X. malinche*. **A**) Principal component analysis of transformed gene abundance counts from ovary RNAseq data from *X. birchmanni* (pink), *X. malinche* (blue), and *X. cortezi* (yellow). For each species, samples included ovarian tissue from both early- and late-stage pregnancies. For *X. birchmanni* and *X. malinche,* two wild-caught samples were also analyzed (denoted by stars and labels). Samples cluster by species, and for *X. malinche*, by developmental stage. Colored envelops show the space in PC1 and PC2 occupied by all samples of a given species. **B**) Prolactin receptor (*prlr*) and releaser (*prlhr*) genes are upregulated in *X. malinche* ovaries, regardless of pregnancy status or sample origin. For each species, mean normalized read count ±2 standard error is shown for early-pregnancy samples on the left (denoted with diamonds) and late-pregnancy samples on the right (denoted with circles). Semi-transparent points show data from each individual (stars for wild-collected). **C)** Expression heatmap of genes from acyl- and acetyl-CoA biosynthesis and vasculature development GO pathways that were enriched in the “darkorange2” WGCNA cluster, which was significantly associated with late pregnancy *X. malinche* ovaries. Sample IDs appear on the x-axis, gene annotations appear on the y-axis. The blue to yellow color bar indicates the difference in expression, with yellow colors indicating greater expression than average and blue colors indicating lower expression than average. **D**) Immunofluorescence demonstrates the presence of the prolactin protein (magenta) in sections from both pregnant *X. birchmanni* females (i.-iii.) and *X. malinche* (iv.-vi.). As expected, prolactin positive cells are identified throughout the developing embryo in mesenchymal and neuronal tissues (such as the eye, i., iv.). Both species also express prolactin in the maternal membranes surrounding the ovary and embryos. Yellow boxes identify the area highlighted in the higher magnification images. iii. and vi. show prolactin signal only, without DAPI counterstain.

Given that late-stage embryos are larger in *X. malinche* than *X. birchmanni*, we tested whether higher genome-wide *X. malinche* ancestry in naturally occurring hybrids predicts larger embryo size. We sampled pregnant mothers carrying stage ≥ 20 broods from the Calnali Low natural hybrid population, since embryo size diverges between species after stage 15. After accounting for stage and brood ID, we found that proportion of *X. malinche* ancestry in the mother’s genome and its interaction with stage are strongly positively associated with embryo dry weight (L-ratio=61.6, p-value<0.0001, R=0.37; Supp. Fig. S6A). Unexpectedly, we find a significant negative relationship between the mother’s ancestry and ovary dry weight (F-value=2.90, p-value=0.02), suggesting that greater *X. birchmanni* ancestry in a hybrid genome predicts higher ovary dry weight.

### X. malinche and X. birchmanni ovaries do not have clear morphological differences

To explore potential differences in ovarian structures between species, we compared histological slides of sectioned ovaries. In highly matrotrophic livebearing fishes, nutrients are delivered to developing embryos through the maternal follicle, which is thicker and highly vascularized (Kwan et al. 2015; Guernsey et al. 2020; Ponce de León and Uribe 2021). Visually, we did not observe consistent differences in maternal follicle thickness, amount of vasculature, or overlap of maternal and embryonic membranes between sections of ovaries carrying unfertilized eggs or late-stage embryos from *X. malinche* and *X. birchmanni* (Fig. 2D; Supplementary Information 6). Given that Matrotrophy Indices for *X. malinche* and *X. birchmanni* both fall below 1, clear morphological differences in reproductive morphology may not be expected (unlike those observed in other studies; e.g. Guernsey et al. 2020).

We also sought to determine whether ovarian follicular tissue is maternal or embryonic in origin (see Supplementary Information 7). Taking advantage of gene expression data from late pregnancy *X. malinche* mothers carrying *X. malinche x X. birchmanni* F_1_ broods, we found that the *X. malinche* allele alone was expressed at nearly all ancestry informative sites, indicating that the ovarian follicle is maternally-derived (Supp. Fig. S7).

### Patterns of gene expression in X. malinche, X. birchmanni, and X. cortezi ovaries

To gain insight into the mechanisms underlying differences in reproductive strategy between these closely related species, we compared gene expression patterns in ovarian tissue from *X. malinche, X. birchmanni,* and *X. cortezi* from “early” pregnancy (fully-yolked unfertilized or stage <5) and “late” pregnancy (25-45; Table S15). Principal component analysis shows that PC1 explains about 31% of the variation in ovary expression and correlates with species while PC2 explains about 17% of total variation and correlates with species and rearing-environment (Fig. 3A). Within *X. malinche*, expression patterns in ovarian tissues separate along PC1 by the developmental stage of the embryos they contained, while in *X. birchmanni* and *X. cortezi* they do not (Fig. 3A). Of the genes expressed in the ovaries of both species (adjusted p-value<0.05), *X. malinche* differs from *X. birchmanni* in expression for 13% (2,202) and 28% (4,934) of genes in early and late pregnancy, respectively, and from *X. cortezi* for 45% (7,957) and 32% (5,639) of genes, respectively. Notably, we see that more genes respond to embryonic developmental stage in *X. malinche* ovaries (310) compared to *X. birchmanni* (63) and *X. cortezi* (130) ovaries. Although we do not focus on them in the main text, we performed similar analyses of genes that were differentially expressed in developing embryos across the three species (Supplementary Information 8).

Several genes that are specifically involved in pregnancy and nutrient provisioning in placental taxa, including mammals and matrotrophic poeciliids, are significantly upregulated in *X. malinche* compared to its close relatives. Notably, we find that the prolactin signaling pathway, which has been implicated in the evolution of matrotrophy in livebearers (Guernsey et al. 2020), is strongly upregulated in *X. malinche* ovaries. Prolactin receptor and releaser genes are highly expressed in *X. malinche* ovaries in both early and late pregnancy compared to *X. birchmanni* and *X. cortezi* ovaries (adjusted p-value<0.001 for all comparisons; Fig. 3B). Prolactin receptors (*prlr*) bind the hormone prolactin and are thought to be directly involved in trophoblast invasion for nutrient uptake in mice (Stefanoska et al. 2013). Prolactin-releasing peptide receptor (*prlhr*) regulates prolactin expression. Interestingly, *prlhr* is also upregulated in late pregnancy *X. birchmanni* ovaries (adjusted p-value<0.001), while we did not detect expression of this gene in *X. cortezi* ovaries (Fig. 3B).

Though there is transcription of prolactin receiving and releasing machinery in late-pregnancy *X. malinche* and *X. birchmanni* ovaries, we do not see expression of prolactin itself based on RNAseq data (as expected since prolactin is a hormone translated in the pituitary gland). To evaluate whether prolactin protein appears in *X. malinche* and *X. birchmanni* ovaries, we performed immunohistochemistry on sections of late pregnancy ovaries for both species (Fig. 3C). We find that prolactin protein is present in both *X. malinche* and *X. birchmanni* maternal ovarian follicle and embryonic tissues. Additionally, as expected, prolactin is present in retinal and mesenchymal tissue of embryos in both species (Fig. 3C).

To explore biological pathways potentially involved in nutrient transfer and provisioning in *X. malinche*, we first looked for enriched Gene Ontology pathways in genes that were significantly differentially expressed between *X. malinche* and *X. birchmanni* ovaries. Notably artery development is enriched, and the genes in this category are upregulated in *X. malinche* ovaries (p-value < 0.002; Supp. Fig. S8). We also see enrichment of several potentially relevant metabolism and nutrient transport pathways, including acyl- and acetyl-CoA metabolism in late-stage *X. malinche* ovaries (Table S16).

We also looked at groups of co-expressed genes and asked which of these groups are significantly associated with late-stage pregnancy in *X. malinche* (see Supp. Fig. S10 for module-trait correlation matrix). Using WGCNA, we identified 17 ovary co-expression gene clusters significantly associated with *X. malinche* ancestry (p-value<0.05), 11 of which were associated with late-pregnancy *X. malinche* ovaries. Gene Ontology enrichment of these clusters revealed that the ‘darkorange2’ cluster was highly enriched for acyl- and acetyl-CoA biosynthesis and metabolic pathways, as well as vascular growth and signaling pathways, like artery development and vasculogenesis (Fig. 3D, Supp. Fig. S9). Many of the represented acyl- and acetyl-CoA metabolism genes are strongly upregulated or downregulated in *X. malinche*, particularly in late pregnancy, compared to *X. birchmanni* and *X. cortezi* ovaries (Fig. 3D, Supp. Fig. S9). In mammals, fatty acids like acyl-CoA are an important source of energy for both placental and fetal growth during gestation and are transferred through the placenta to the fetus during mid-to later-stages of pregnancy (Chavan-Gautam et al. 2018). The ‘darkorange’ cluster was enriched in amino acid biosynthesis and metabolism pathways, and the ‘violet’ cluster was enriched for intracellular and transmembrane signaling pathways.

### Testing the potential role of resource availability in driving the evolution of large offspring size

We tested the response of offspring of both species to two ecological factors that differ between the *X. malinche* and *X. birchmanni* environments. We found that the larger offspring of *X. malinche* did not have improved minimum thermal tolerance compared to *X. birchmanni* fry (Supplementary Information 11). We thus turned to investigating another ecological factor that correlates with temperature: food availability. Cooler, headwater habitats are generally less resource-rich than warmer downstream habitat (Reznick 1982*b*). Given low winter temperatures observed in *X. malinche* populations (Supp. Fig. S11; Payne et al. 2022), we asked whether cold temperatures (and presumably low primary production and food availability associated with such temperatures) were linked to seasonal breeding in *X. malinche* populations and starvation tolerance in newborn fry.

Water temperatures at *X. malinche* and *X. birchmanni* sites are coolest from November to March with peak lows in January and February and are warmest from April to October with peak highs in May and June (Supp. Fig. S11). The shifts in temperature roughly correspond with the rainy season and river flooding, which typically starts late June and ends in October. We collected mature females from natural populations of both *X. birchmanni* and *X. malinche* across seasons. We found that while many *X. birchmanni* females were reproducing regardless of sampling month or season (the lowest pregnancy rate observed was 34% in February), <20% of *X. malinche* mature females were pregnant from February – August, while 80% were pregnant in May (Supp. Fig. S12; Table S22).

To evaluate how relative female condition may vary seasonally, we compared total lipids from wild nonpregnant Chicayotla *X. malinche* and Coachuilco *X. birchmanni* females and found temporal differences in fat content between species. In February, before the *X. malinche* breeding season, we found no significant difference in percent fat content between *X. malinche* and *X. birchmanni* females (t-value=1.64, p-value=0.12; Fig. 4A). However, in September, after the *X. malinche* breeding season, *X. birchmanni* females had significantly higher fat content than *X. malinche* (t-value=-12.45, p-value=1.30×10^-6^), consistent with limited resource availability in *X. malinche* habitat.

**Figure 4.**
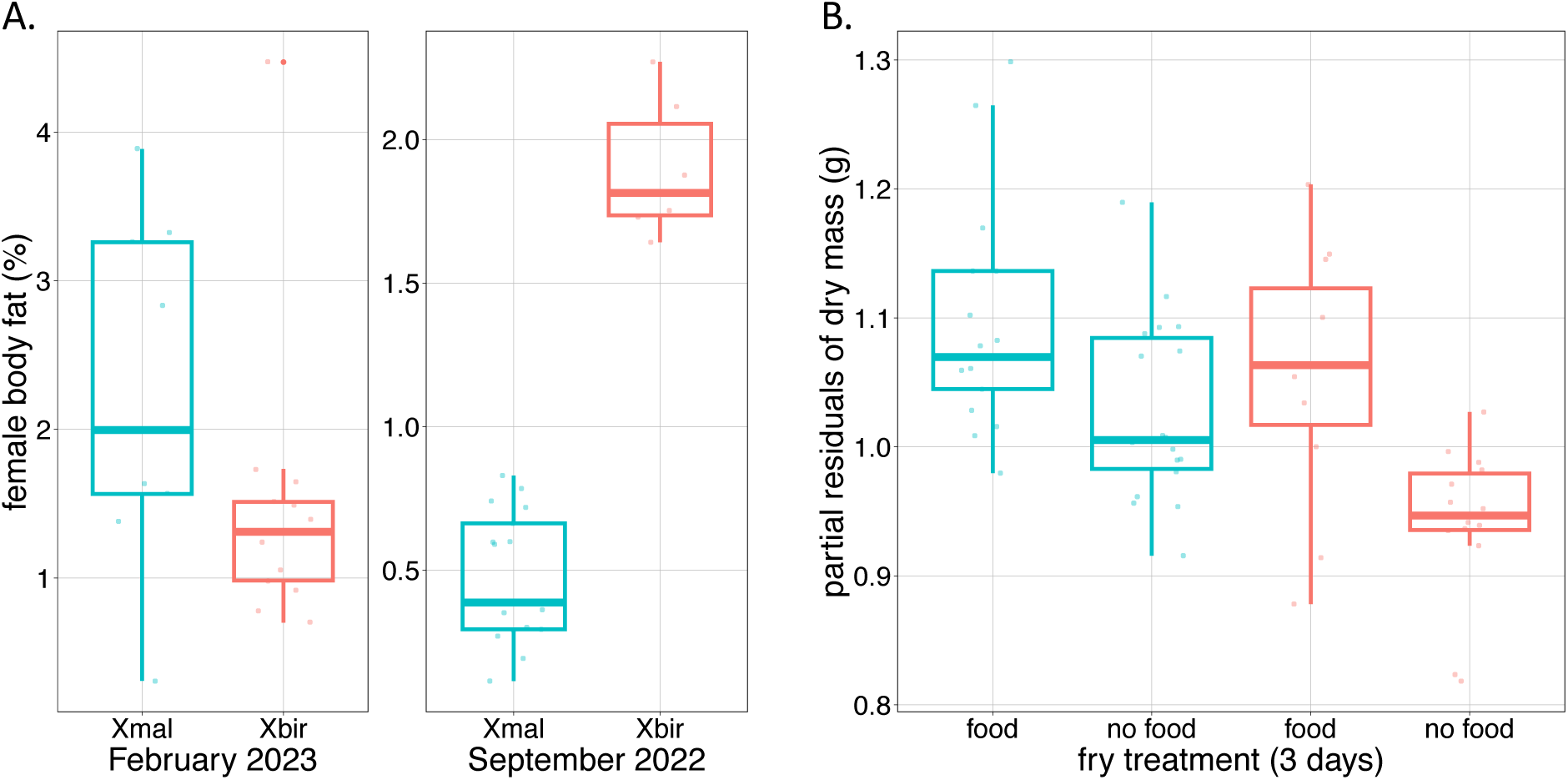
Comparisons of responses to food availability in *X. malinche* and *X. birchmanni*. **A**) Boxplots showing fat content of non-pregnant *X. malinche* and *X. birchmanni* females collected from wild populations in different seasons. *X. malinche* females have much lower fat content before their breeding season than *X. birchmanni*. Note that fat content is not directly comparable across the two collections given that samples were processed in different batches. **B**) Results of starvation experiments in newborn *X. birchmanni* and *X. malinche* fry. *X. birchmanni* (pink) and *X. malinche* (blue) did not significantly differ in weight three days after birth when both were fed, indicating that *X. birchmanni* fry gain weight rapidly after birth (see Fig. 1A). However, in starvation conditions, *X. birchmanni* fry weighed significantly less than *X. malinche* fry, suggesting that *X. malinche* fry have improved tolerance to starvation.

To directly compare how newborn fry of the two species respond to reduced food availability, we exposed *X. malinche* and *X. birchmanni* newborn fry to abundant-food and no-food conditions and compared standard length, dry weight, and fat content between treatments and species. Surprisingly, after three days of feeding, *X. malinche* and *X. birchmanni* fry were approximately the same standard length and dry weight (ANOVA and Tukey post-hoc test p-value=0.76 and 0.91, respectively). Compared to their initial standard lengths, *X. birchmanni* fry grew 4× more rapidly (∼0.32mm/day) than *X. malinche* fry (∼0.08mm/day) (Supp. Fig. 13A). However, while both *X. malinche* and *X. birchmanni* unfed newborn fry weighed less than their fed counterparts, we found that on average food deprivation for three days impacted *X. malinche* fry weight less than *X. birchmanni* fry, suggesting that they may be more resilient to starvation immediately after birth (Fig. 4B). We found no significant difference in total body fat content between species for either treatment (Supp. Fig. S13B).

### Differences in viability by cross direction

In generating F_1_ crosses for this and other projects, we have found a clear difference in the frequency of premature birth and stillbirth in crosses between *X. malinche* mothers and *X. birchmanni* fathers compared to crosses between *X. birchmanni* mothers and *X. malinche* fathers. Given that these crosses are genetically identical except for the Y chromosome and mitochondrial haplotype (Moran et al. 2021), this raises the possibility that maternal effects may impact survival of the F_1_ fry.

We found high rates of premature birth in crosses between *X. birchmanni* mothers and *X. malinche* fathers (Table S23). Of 25 crosses generated by artificial insemination between 2011-2022, only eight F_1_ fry survived (out of 64 born). In five of these crosses, the mother also suffered mortality. In sixteen out of the 25 crosses, mothers gave birth to morphologically premature fry (Supp. Fig. S14), which have a low probability of survival (∼98% mortality, (Moran et al. 2021). In the ten crosses where fry were born at term, 50% contained stillborn fry, versus 7% for premature broods (Fisher’s exact p-value=0.04). Maternal mortality was observed in 30% of these full-term crosses, versus 13% of crosses where fry were born prematurely (though this difference was not significant; Fisher’s exact test p-value=0.1). For cases where we developmentally staged premature fry, individuals tended to be around stage 40.

By contrast, we have never documented a premature birth, stillbirth, or maternal mortality for *X. malinche* mothers. To date we have generated >190 adult F_1_ hybrids and 1,253 adult F_2_ hybrids from intercrossing these individuals. Premature birth has also not been observed in F_2_ crosses between these F_1_ individuals.

## Discussion

Offspring size is a complex trait that impacts both parent and offspring fitness. Compared with its sister species, *X. birchmanni*, and other *Xiphophorus* species, *X. malinche* produce exceptionally large fry. Though offspring size has been shown to be sensitive to ecological conditions and the maternal environment in livebearers (Reznick 1982*a*; Reznick and Yang 1993; Reznick et al. 1996*b*; Bashey 2006*b*, 2008), we see this size difference in the offspring of lab-raised individuals, and estimate that the broad-sense heritability of offspring size attributable to ancestry is 0.25. Moreover, the proportion of the genome derived from *X. malinche* impacts offspring size in natural hybrid populations.

Larger offspring size generally increases the probability of survival to adulthood across a variety of species (Brockelman 1975; McGurk 1986; Tessier and Consolatti 1989; Einum and Fleming 2000; Marshall and Keough 2007; Jørgensen et al. 2011; Rollinson and Hutchings 2013; Bowen et al. 2015). Given that *X. malinche* live in unique environments compared to other species in the genus, we predicted that larger offspring size may have evolved in response to ecological pressures – specifically, limited nutrient availability. Sampling for this study revealed that while *X. birchmanni* breed across seasons, *X. malinche* breed more seasonally, with peak productivity in the warmest months. Notably, before the *X. malinche* breeding season, we find that nonpregnant *X. malinche* and *X. birchmanni* females have comparable body fat content. However, after the *X. malinche* breeding season, *X. malinche* have significantly lower body fat content than *X. birchmanni* females, hinting that resources are generally more limited in *X. malinche* habitat.

Past work in guppies has shown that under limited food conditions, mothers produce larger offspring (Reznick 1982*b*; Reznick and Yang 1993; Reznick et al. 1996*b*; Bashey 2006*b*), which presumably have a fitness advantage in lower resource environments (Pettersen et al. 2015, 2022). We directly tested offspring performance in food-depleted conditions in a controlled lab experiment. We find that starved newborn *X. malinche* retain more weight than newborn *X. birchmanni*. This suggests that *X. malinche* fry may be more resilient to a low resource environment, consistent with resource limitation (or high competition) as a driver of the evolution of large offspring in *X. malinche* (Bashey 2006*b*, 2008). Notably, in scenarios where resources were not limited, newborn *X. birchmanni* fry grew more rapidly than *X. malinche* and matched their size within days. This pattern could hint that there is an advantage to being large early, consistent with previous work in other taxa (Chambers et al. 1989; Gliwicz and Guisande 1992; Bashey 2008).

There are two mechanisms through which livebearing fish can produce larger fry. First, mothers may produce larger eggs. Second, mothers may continue to provision nutrients to offspring over embryonic development (note that matrotrophy does not predictably lead to large offspring at birth across taxa; (Furness et al. 2021). We evaluated evidence for both mechanisms in *X. malinche*. We measured dry weights for embryos across ten embryonic stages ranging from unfertilized fully-yolked eggs to fully developed juveniles for *X. malinche, X. birchmanni,* and their close relative, *X. cortezi*. Notably, unfertilized fully-yolked eggs and early stage embryos are similar in size between *X. malinche* and *X. birchmanni* (Fig. 2). However, *X. malinche* and *X. birchmanni* embryos diverge in size around stage 20 of embryonic development. While *X. birchmanni* embryos undergo an overall decline in dry weight after stage 15, *X. malinche* embryos maintain roughly the same weight over development. We summarize this difference in developmental profile with the first estimates of Matrotrophy Index for *X. malinche* (0.98) and *X. birchmanni* (0.66). This estimate for *X. malinche* is consistent with *X. malinche* embryos receiving some nutrients from their mother over development, contributing to larger offspring size at birth in *X. malinche*. It is notable that although *X. birchmanni* do not appear to provision nutrients during development, *X. birchmanni* also has fry that are much larger at birth than other *Xiphophorus* species. We were unable to collect sufficient samples to accurately quantify Matrotrophy Index from these species but we predict that they may produce smaller eggs than *X. birchmanni*, and indeed the few samples we have are consistent with this prediction (Fig. S17).

Research on highly matrotrophic species (e.g. with Matrotrophy Indices>40) has shown that maternal and embryonic tissues interact to form an interface for nutrient transfer, with some species forming placental structures of varying complexity (Guernsey et al. 2020). To look for evidence of a more complex maternal-offspring interface in *X. malinche*, we quantified characteristics of the maternal-embryo interface in histological sections of late-stage pregnancy in *X. malinche* and *X. birchmanni* ovaries. In highly matrotrophic poeciliids, embryonic and maternal blood vessels range from apposed to fused (Ponce de León and Uribe 2021) and the ovarian follicular epithelium is dense and maximizes surface area contact with embryos for nutrient transfer (Kwan et al. 2015). We did not see consistent differences in ovarian follicle thickness or vascularization between stained sections of late-stage *X. malinche* and *X. birchmanni* ovaries. However, we found it difficult to compare these traits between 2-dimensional slices of ovary. Because post-fertilization nutrient provisioning in *X. malinche* arose recently and is less substantial than that observed in other species, we might predict differences in ovary morphology between these species to be subtle. Future work should focus on obtaining 3-dimensional images suitable for quantitative comparisons of the whole ovary.

To further investigate differences in maternal nutrient provisioning at the molecular level, we quantified gene expression patterns in early- and late-stage ovarian tissue from pregnant *X. malinche* and *X. birchmanni*. The observed expression patterns raise multiple possible mechanisms through which post-fertilization nutrient provisioning may be occurring in *X. malinche*. Receptor and releaser genes for prolactin, a pituitary hormone important during pregnancy in mammals and in matrotrophic poecilids (Menzies et al. 2011; Guernsey et al. 2020), are highly expressed in the *X. malinche* ovarian follicle compared to *X. birchmanni* (and *X. cortezi*) follicles both before and during pregnancy. Combined with evidence that *X. malinche* mothers appear to provision their embryos over development to a greater extent than *X. birchmanni* mothers, this result suggests that prolactin may be a key candidate for the evolution of matrotrophy in livebearers (as proposed by Guernsey et al. 2020). Immunostains of prolactin in sections of late-stage pregnant *X. malinche* and *X. birchmanni* ovaries confirms that prolactin is present in the ovarian follicle of both species. Given these results, we predict that the striking differences in abundance of prolactin releasing and receiving machinery that we observe at the gene expression level (Fig. 3) may regulate the downstream transfer and function of prolactin in the two species, which includes nutrient transport.

While offspring almost universally benefit from being larger at birth, producing large offspring can be costly for parents (Einum and Fleming 2000; Walker et al. 2008; Ljungström et al. 2016) and constrained by environmental conditions (Reznick and Endler 1982; Hutchings 1991; Janzen et al. 2000; Allen et al. 2008; Marshall and Keough 2008; Pettersen et al. 2022). These pressures can result in parent-offspring conflict in the evolution of offspring size (Trivers 1974; Crespi and Semeniuk 2004; Moore 2012). Similarly, if males and females differ in their level of resource investment in offspring, conflict between the maternal and paternal genomes can drive similar dynamics (Moore and Haig 1991), driving evolutionary arms races between paternally contributed factors that increase offspring growth and maternally contributed factors that suppress them (Moore and Haig 1991; Barlow and Bartolomei 2014). In hybrids between species, misregulation of these interactions can result in larger or smaller offspring than expected (i.e. depending on the cross direction) and impact viability of hybrid offspring (Shi et al. 2004; Brekke & Good 2014), including in livebearing fish (Schrader et al. 2013). Given these findings and the large difference in *X. malinche* and *X. birchmanni* embryos during development, we were curious to understand whether this trait impacts reproductive isolation between these species, which naturally hybridize in the wild (Culumber et al. 2011). In the lab, *X. malinche* mothers and *X. birchmanni* fathers successfully produce viable F_1_ offspring and subsequent early-generation crosses, whereas the reverse cross is typically aborted late in pregnancy (by stage 40). Asymmetric reciprocal F_1_ cross viability has been shown to stem from differences in nutrient provisioning over development in other poeciliid species (Turcotte et al. 2008; Schrader et al. 2013). However, this case is unique in that F_1_s are inviable when the mother is lecithotrophic and therefore completely controls nutrient provisions (i.e. the matrotrophic paternal genome cannot affect nutrient allotment). We initially hypothesized that differences in provisioning strategy might result in embryos that grow too large too quickly for *X. birchmanni* mothers to sustain. However, we found it difficult to collect stage matched offspring in F_1_ hybrid crosses with *X. birchmanni* mothers, resulting in a low sample size or this comparison (Fig. 2D). While initial results do not suggest size differences compared to stage matched *X. birchmanni* embryos, rigorously testing this hypothesis is an important direction for future work. Alternatively, the energetic demands of F_1_ embryos may outpace the nutrients stored in yolk from *X. birchmanni* mothers early on, which could lead to malnourishment. Consistent with this hypothesis, we find that many offspring from this cross are stillborn and the majority are born prematurely.

Our data also shows that offspring size is a complex trait, and the size of any individual offspring is impacted by many variables, including developmental stage, mother size, and mother environment. Accounting for all of these effects make this trait difficult to characterize, complicating comparisons of offspring size and embryonic development profiles between livebearing species. Recently, Skalkos et al. (2023) reviewed current limitations in how matrotrophy is studied for teleost fish. Our findings underscore some of their points, including that direct measures of early- and late-stage embryo weight, rather than predictive measures using regression-based approaches, should be used to estimate Matrotrophy Index where possible (since stages are morphological categories that do not linearly track developmental time). Moreover, if provisioning varies over development (as hinted at by some of our data), more nuanced analysis methods may be more appropriate. Future work should consider these factors. We note that in our work using lab populations served as a powerful resource for confirming patterns inferred from wild populations (Fig. 2), as they allowed us to control for many of the environmental variables that impact wild collections.

Using a combination of morphometric and molecular data, we document immense variation in offspring size across *Xiphophorus* species and provide the first evidence of incipient matrotrophy and post-fertilization nutrient provisioning mechanism in the *Xiphophorus*. Importantly, because *X. malinche* and *X. birchmanni* are recently diverged (∼250,000 generations ago), this is an example of recent shift in reproductive strategy. Parent of origin incompatibility phenotypes associated with spontaneous abortion late in development hint that this difference in reproductive strategy could contribute to the diverse reproductive barriers present in these young sister species.

## Data Availability

All data is publicly available. Raw sequences are available under NCBI BioProject PRJNAXXXX. All code and data files can be found in the accompanying GitHub repository at https://github.com/cypayne/swordtail-offspring-size.

## Acknowledgements

We thank Moi Exposito-Alonso and members of the Schumer lab for helpful feedback on previous versions of this manuscript. We are grateful to the Mexican federal government for permission to collect samples. We thank Stanford University and the Stanford Research Computing Center for providing computational support for this project. This study was supported by a Society for Integrative & Comparative Biology GIAR grant to CYP, an American Society of Naturalists Student Research Award to CYP, and a Human Frontiers in Science Programme grant (RGY0081), HHMI Freeman-Hrabowski award, Pew Biomedical Scholars, and Searle Scholars Award to MS.

