## Supplementary Information for "Recent evolution of large offspring size and post-fertilization nutrient provisioning in swordtails"

**Supplementary Materials**

*Supplementary Materials 1. Life history traits in X. malinche and X. birchmanni*

Since mother body size correlates with offspring size in many species of poeciliids, we first tested whether *X. malinche* and *X. birchmanni* females differ in adult size when grown in common conditions. *X. birchmanni* males are larger bodied than *X. malinche* males (Preising et al., 2022), but this trend goes in the opposite direction of our expectation given newborn size.

To test this, we used standard length measurements from females grown in outdoor mesocosms in common conditions (N=35 *X. birchmanni;* N=44 *X. malinche*). While *X. malinche* females were slightly larger on average (4.04 cm versus 3.94 cm in *X. birchmanni*), between tank variation in size exceeded between species variation in size (Supp. Fig. S4). Nonetheless, wherever possible, we included mother standard length in our analyses of differences in embryo or offspring size (see Methods).

We also wanted to evaluate the relationship between brood size and species and mother size. To do so, we applied a linear model approach implemented in R using our large dataset of mothers and embryos collected from natural *X. malinche* and *X. birchmanni* populations. For each brood, we calculated the average weight of offspring by stage and summarized the number of offspring per brood. After model selection, we analyzed average embryo dry weight as a function of species, stage, maternal standard length, and collection season. Pregnant mothers collected from the wild for this study ranged in standard length from 3.4-5.7cm for *X. malinche* and 2.8-5.2cm for *X. birchmanni*.

As expected, given reports in numerous poecilid species, we saw a positive effect of mother standard length on offspring number, with larger mothers having more offspring per brood (F-value=95.4; p-value=$6.4 \times 10^{-16}$). We also found a significant effect of mother standard length on the average weight of the embryos in a brood (F-value=6.7; p-value=0.01).

We also wanted to determine whether the number of offspring per brood systematically differed between species. Surprisingly, we found that *X. malinche* on average had slightly larger broods (on average 17 fry, compared to 13 in *X. birchmanni*). This difference remained significant after accounting for mother standard length and season (F-value=4.7; p-value=0.03). Thus, *X. malinche* produce slightly more offspring per brood on average in addition to larger offspring on average. However, due to seasonality in breeding in *X. malinche* (see main text), we do not expect that the total reproductive output of *X. malinche* exceeds that of *X. birchmanni*.

*Supplementary Materials 2. Estimating broad-sense heritability of offspring size*

We were interested in obtaining an estimate of heritability of offspring size, particularly the heritability that is driven by *X. malinche* ancestry or *X. birchmanni* ancestry. Using lab-reared fry of the two species and fry from $F_1$ and $F_2$ hybrid crosses, we obtained an estimate of the broad-sense heritability ($H^2$) of standard length at birth attributable to species-level differences using the formula:

$$H^2 = \frac{(\sigma_{F2}^2 - \sigma_{env}^2)}{\sigma_{F2}^2}$$

where $\sigma_{F2}^2$ is the within population variance for the $F_2$ population and $\sigma_{env}^2$ is the variance attributed to the environment (as in Fraser, 2020). Model selection with the R stats::step function indicated that season has a significant effect on fry length even in the lab environment. Collection dates were binned into two seasonal categories describing water temperature, which matches that of wild populations (Supp. Fig. S11): the "warm" season includes April-October months and the "cold" season includes November-March months. Therefore, we accounted for the effect of season by calculating partial residuals for the following mixed linear model and included brood ID as a random effect, treating the partial residuals as the trait of interest: fry standard length ~ species + season + (1|brood ID). Partial residuals can be found in Table S24. Since $F_1$ hybrids are all heterozygous for *X. birchmanni* and *X. malinche* ancestry across the genome, they can be used, along with the variance observed within the parental species, to estimate the variation in offspring size attributable to the environment (or segregating variation within the parental species). We calculate this value (0.45) with Wright's weighted average of within-population variance, using the formula (following (Fraser, 2020):

$$\sigma_{env}^2 = \frac{\sigma_{p1}^2}{4} + \frac{\sigma_{p2}^2}{4} + \frac{\sigma_{F1}^2}{2}$$

*Supplementary Materials 3. Matrotrophy estimates and embryo size*

Since we opportunistically collected pregnant females from wild populations to generate profiles of embryonic growth during development, we were limited by the number of pregnant females we were able to collect and the developmental stage of embryos at the time of collection.

We were interested in estimating Matrotrophy Index for *X. cortezi*, which is the sister group to the *X. birchmanni* and *X. malinche* clade. Data for *X. cortezi* will be especially important for building a model of the evolution of matrotrophy in the genus since offspring size at birth is very small. Based on a small sample size of four broods, the average size of fully-yolked eggs in *X. cortezi* is smaller than the average *X. birchmanni* and *X. malinche* egg size, though not significantly after accounting for brood size and mother standard length (adjusted p-value>0.40; Fig. S17). Unfortunately, we only collected one stage 50 brood from *X. cortezi*, making estimates of Matrotrophy Index unreliable.

Given that mother standard length and season impact embryo dry weights, we calculated Matrotrophy Index for *X. birchmanni* and *X. malinche* using partial residuals of averaged dry weights accounting for these covariates and report those values in the main text (0.66 and 0.98, respectively). We also calculated Matrotrophy Index using the raw averaged dry weights for stage 50 and stage 10 embryos, which yielded a Matrotrophy Index of 0.66 for *X. birchmanni* and of 0.95 for *X. malinche*. Thus, regardless of the approach we use to quantify Matrotrophy Index, our estimate for *X. malinche* exceeds our estimate for *X. birchmanni*.

In an effort to evaluate whether the difference in Matrotrophy Index between *X. malinche* and *X. birchmanni* was significant, we performed bootstrapping of the data used in the Matrotrophy Index calculation. For each species, we resampled the values for average embryo dry weight of broods at stage 10 and stage 50 brood with replacement and recalculated Matrotrophy Index as described above 1000 times. This gave us a 95% confidence interval of 0.656-0.659 for *X. birchmanni* and 0.955-0.965 for *X. malinche*. This indicates that the difference in Matrotrophy Index between species is significant.

*Supplementary Materials 4. Genome-wide ancestry and mitotype is associated with embryo size in natural hybrids*

As described in the main text, we collected paired mother and embryo data from 35 pregnant females from the Calnali Low hybrid population. We have worked extensively on this population in past work for admixture mapping (Moran et al., 2021). To estimate maternal ancestry across the genome, we relied on local ancestry inference approaches developed by our group (Schumer et al. 2020). Specifically, we generated ~1X whole genome sequence data using a tagmentation-based library preparation protocol (following Moran et al., 2021) and performed

HMM-based local ancestry inference using the *ancestryinfer* pipeline. Based on previous analyses of the Calnali Low hybrid population, we used 40 generations as the prior for time since initial admixture and 0.5 for the prior expectation for admixture proportion. Past work has indicated that local ancestry inference in *X. birchmanni* x *X. malinche* hybrids is highly accurate due to the high density of fixed ancestry informative sites and the recency of admixture in these populations (Schumer et al. 2020).

For each individual, *ancestryinfer* outputs posterior probabilities of each ancestry state at ancestry informative sites along the genome (i.e. homozygous *X. birchmanni*, heterozygous, or homozygous *X. malinche*). We chose a hard-call threshold of posterior probability of 0.9 and converted posterior probabilities at sites that exceeded that threshold to the highest probability ancestry state. For sites where no ancestry state was supported at ≥0.9 posterior probability, we converted the genotype at that site to NA. We used these hard-calls to estimate the proportion of the genome derived from *X. malinche* and *X. birchmanni* in each hybrid individual, which allowed us to analyze the correlations between embryo size and maternal admixture proportion, as described in the main text.

Previous work from our group has shown that hybrid incompatibilities between nuclearly encoded genes and mitochondrially encoded genes can impact embryonic development in hybrids (Moran et al., 2021). In particular, individuals with a *X. malinche* mitochondria and homozygous *X. birchmanni* ancestry at the gene *ndufs5* experience stunted embryonic development. For our analysis of natural hybrids, it is plausible that some embryos with *X. malinche* mitochondria may be smaller because of the direct effects of this hybrid incompatibility. However, since mitochondrial ancestry in hybrids from this population is correlated with genome-wide ancestry (Spearman's $\rho$=0.69), we might reasonably expect that embryos with *X. malinche* mitochondria would be larger.

Consistent with our expectation based on maternal ancestry alone, we find that the *X. malinche* mitotype predicts larger embryo size (L-ratio=11.8, p-value=0.0006; Supp. Fig. S6B). This suggests that the hybrid incompatibility between the *X. malinche* mitochondria and *X. birchmanni* ancestry at *ndufs5* is not having a major impact on our results from natural hybrids. We found no relationship between maternal mitochondrial ancestry and ovarian tissue dry weight (F-value=0.028, p-value=0.87).

*Supplementary Materials 5. Results of within species crosses*

We also made crosses between populations of *X. malinche* to investigate embryo size variation in within species crosses. *X. malinche* has low genetic diversity and past work has indicated that this lineage has experienced a sustained bottleneck and accumulated an excess of nonsynonymous mutations compared to *X. birchmanni* (Schumer et al., 2018). We crossed Chicayotla *X. malinche* males with females from the allopatric Tetipanchalco *X. malinche* population, which occur in tributaries in different rivers (Río Xontla and Río Claro, respectively). We lack access to similar populations for a within species cross for *X. birchmanni.* We measured dry weight of embryos from two successful intrapopulation crosses, which were at stages 25 and 35, and dry weight of ovarian tissue as described above. We subsampled within population crosses for broods with embryos between stages 25 and 35, and statistically compared embryo dry weight, correcting for mother standard length and brood size as selected covariates and brood ID as a random effect with R lme4::lmer, between the within- and between-population crosses using an ANOVA and Tukey post-hoc test with R emmeans::emmeans (see Table S11 for embryo size data and Table S12 for statistical comparison of means). Partial residuals of embryo dry weight were calculated and plotted with R visreg::visreg (Fig. S15).

Stage 25 to 35 Chicayotla embryos trend towards being larger than Tetipanchalco embryos (though not significantly, p-value=0.06). Stage 25 to 35 embryos from Tetipanchalco mothers and Chicayotla fathers were intermediate and did not significantly differ in size from within population cross embryos (Table S12).

*Supplementary Materials 6. Morphological comparison of maternal-embryo interface*

We attempted to quantitatively evaluate whether there were differences in ovary morphology between species using the digital scans of H&E stained sections of pre-fertilization (stage 0) and late-stage pregnancy (stage 35-40) *X. malinche* and *X. birchmanni* ovaries (see *Multifactorial artificial insemination crosses* methods in the main text). In QuPath v0.5.0, we measured the length of overlap of maternal and embryonic vasculature, maternal follicle thickness, and embryonic pericardial sac thickness. We used a publicly available script (https://gist.github.com/petebankhead/bc4f1bc8c3e32aba9f19e0d85c6753b4) to randomly place 50 rectangles (1000x1000 pixels) across each digital slide. Rectangles that did not include either the maternal follicle or the pericardial sac and/or were clearly damaged were excluded from analysis. For each embryo (typically 1-2) within a rectangle we measured the thickness of the embryonic pericardial sac and the thickness of the nearest maternal follicle to the nearest 0.0001 µm at its thickest point. When the maternal follicle and embryonic pericardial sac were apposed (i.e. with $\leq$ 5 µm gap) within the rectangle, this was labeled an overlap and its length was measured until the gap between tissues exceeded 5 µm. Some overlaps extended beyond the rectangle boundaries, but none crossed into a second rectangle.

We note several limitations with our data collection approach. Given that we only sampled histology for one stage 0 and one late-stage ovary from each species, it is not possible to make general statements about ovary morphology. In addition to more biological replicates, we believe that data quality and completeness would be much improved by collecting 3D images of ovaries. Therefore, we chose not make statistical comparisons of overlap length, maternal follicle thickness, or pericardial sac thickness between species or pregnancy stage.

However, we note some general observations from the data we collected here. Total overlap length per embryo may decrease from early to late stage pregnancy for both species. Maximum (i.e. thickest value recorded) and mean maternal follicle thickness per embryo may decrease from early to late stage pregnancy for *X. birchmanni*. Mean embryo pericardial sac thickness per embryo may increase from early to late stage pregnancy for both species. Though we are hesitant to compare between species given the caveats described above, *X. malinche* may maintain similar maternal follicle and embryo pericardial sac thickness across development. Future statistical tests using biological replicates and 3D scans ovaries for each species across developmental stages will allow us to make robust comparisons of ovary morphology.

*Supplementary Materials 7. Determining the genetic origin of tissue spanning the maternal-embryo boundary*

In mammals and some other vertebrates, placental tissues are fetally-derived (Blackburn, 2015; Whittington et al., 2022). To investigate whether tissues spanning the maternal-embryo boundary in *X. malinche* females were mother or offspring derived, we looked at allelic expression in RNAseq data in tissue collected from *X. malinche* mothers pregnant with $F_1$ hybrid offspring. Since offspring are $F_1$ hybrids, equal expression of *X. malinche* and *X. birchmanni* alleles at genes across the transcriptome would indicate that the interface tissue originated from the offspring. Alternately, expression of only *X. malinche* alleles would indicate that the tissue is maternal in origin. To quantify this allelic representation, we took advantage of additional RNAseq data for two *X. malinche* mothers pregnant with $F_1$ offspring (extracted, prepared, and sequenced as described in the Methods) and compared allele-specific expression across the genome using the tool ncASE (https://github.com/YourePrettyGood/ncASE). Briefly, ncASE maps RNAseq reads to both parent reference transcriptomes, calls variants, and filters out sites that show mapping bias to one of the references. We subset read counts for all ancestry informative SNPs in any gene across the transcriptome and evaluated the ratio of *X. malinche* to *X. birchmanni* alleles.

*Supplementary Materials 8. Embryo differential gene expression between species*

In the main text, we describe the results of an experiment where we performed artificial insemination and collected ovarian tissue from *X. malinche* mothers carrying *X. malinche* broods, *X. birchmanni* mothers carrying *X. birchmanni* broods, and *X. cortezi* mothers carrying *X. cortezi* broods. For each species, we collected early and later stage pregnant females (see main text). For each brood of later stage embryos (stage $\geq$ 25), we also collected and prepared an RNAseq library for one developing embryo, as described in the main text. We expect that differences in expression between groups of embryos will be dominated by developmental stage differences (and to a lesser extent, evolved species-level differences). However, this data may also contain some clues about the role of embryonic signaling in nutrient provisioning, so we considered it worthwhile to collect this data.

We performed differential expression analysis for these embryonic samples as described in the main text (under *Differential gene expression and co-expression network analysis*). As expected, PCA analysis shows that individual samples separate by developmental stage and species to a lesser extent (Supp. Fig. S16). Out of a total of 19,176 genes in the *X. birchmanni* reference transcriptome, we find that 18,322 are expressed in at least one of the species' embryos in mid- to late-pregnancy. Notably, only 315 genes are differentially expressed between *X. birchmanni* and *X. malinche* embryos (at an adjusted p-value < 0.05), while *X. cortezi* expression differs for 4,630 and 5,237 genes with each of these species, respectively, though this difference is likely driven, at least in part, by the embryonic stages sampled. We expect that this dataset will be useful to reference as we gain a better understanding of the role of embryo-mother communication during nutrient provisioning and the mechanisms through which it acts.

*Supplementary Materials 9. Co-expression network analysis with WGCNA*

Unsupervised clustering of co-expressed genes can be used to associate interacting genes and biological pathways with a trait of interest. We used the R package WGCNA to evaluate patterns of co-expression in the ovary RNAseq data from *X. birchmanni*, *X. malinche*, and *X. cortezi (Langfelder & Horvath, 2008)*. WGCNA clusters genes by their expression patterns across samples into groups called modules and summarizes overall expression in modules with 'module eigengenes' (i.e. PC1 of the gene expression of all genes in the module). This allows us to correlate traits of interest, like species and developmental stage, with module expression profiles, and identify enriched biological pathways within correlated modules.

To ensure samples had comparable variances, raw gene counts were normalized for library size and size factors (the median ratio of the geometric mean of a gene over all samples) with DESeq2 varianceStabilizingTransformation. Genes with missing data for half the samples or no variance across the 27 samples were dropped (260 dropped, leaving 18,916 for clustering). Using WGCNA's blockwiseModules function, we generated a single-block unsigned gene network with a minimum module size of 20 and a soft-thresholding power of 8, as recommended by the WGCNA documentation for a sample size of 20-30. The resulting network clustered genes by strength of co-expression regardless of the direction of expression correlation (i.e. positive or negative).

We next evaluated correlations between module eigengenes and the interaction between species (*X. malinche, X. birchmanni, X. cortezi*) and pregnancy category (early or late) with the WGCNA corPvalueStudent function. The modules that correlated with any of these variables at Student asymptotic p-value < 0.05 were tested for an enrichment of GO biological pathways (see the *Differential gene expression and co-expression network analysis* methods in the main text).

*Supplementary Materials 10. Immunostaining of prolactin expression in ovarian tissue*

Using unstained slides from individuals sectioned for histology (see Methods section *Multifactorial artificial insemination crosses*), we immunostained late-stage pregnancy ovaries for prolactin. Slides were deparaffinized through two 10-minute washes in xylenes and rehydrated through two 5-minute washes each in a graded ethanol series (100%, 90%, 80%, 70%, 50%, $dH_2O$). Slides were then blocked in serum block (5% normal goat serum; Jackson ImmunoResearch 005-000-121, 1% BSA, 0.01% Triton-X in PBS) for 1 hour at room temperature. Anti-prolactin primary antibody (rabbit, Abcam, EPR19386) was diluted 1:200 in serum block and incubated overnight at 4C. Endogenous peroxidase was then blocked with 3% peroxide in PBS for 10 minutes at room temperature. Biotinylated goat anti-rabbit secondary antibody (Jackson ImmunoResearch 111-065-144) was diluted 1:5000 in serum block and incubated for 1 hour at room temperature. Slides were then incubated with VectaShield Elite ABC reagent (Vector Labs PK-6100) for 30 minutes at room temperature, followed by incubation with TSA-Cy3 (Akoya Biosciences NEL744001KT) for 6 minutes at room temperature and protected from light. Slides were counterstained with DAPI (Thermo Scientific 62248) 1:2000 in PBS for 5 minutes, washed in PBST, and mounted with Prolong Gold Antifade (ThermoScientific P36930) and #1 cover glass (VWR 43893-106) and allowed to harden for 24 hours before imaging. All incubations were performed in a humidified chamber. After blocking step, all incubations were followed by three 10-minute washes in PBST (PBS + 1% Tween-20).

Samples were imaged with a Nikon Ti Eclipse inverted microscope equipped with an ASI MS-2000 motorized linear XY stage, Yokogawa CSU-W1 single disk (50mM pinhole) spinning disk unit, Andor Zyla 4.2 (6.5mM pixel size) scMOS camera, and 10x/0.45 NA or 20X/0.75 NA Nikon PlanApo Lambda air objectives. The final digital resolution of the images was 0.65mm/pixel or 0.325mm/pixel, respectively. Blue and red fluorescence was collected by illuminating the sample with a 405nm or 594nm laser, respectively, in a SPECTRA laser launch, then acquired sequentially through either an ET450/40M (blue) or ET595/50M (red) emission filter. Nikon Elements v4.30.02 was used to acquire the data. Images of entire embryos within the ovary were captured by tile-scanning the tissue with 15% overlap for stitching after manually identifying XY tissue boundaries. Tiles were manually focused in Z across the scan area to generate a focus surface which determined precise z-position during acquisition. Images acquired at a single field of view were also taken at a single z-position. Data was saved in .nd2 format and manually processed in ImageJ.

*Supplementary Materials 11. Measuring cold tolerance with $CT_{min}$ trials*

We were interested in exploring possible physiological impacts of differences in offspring size, especially those that interact with the ecological environment. One major environmental difference between *X. birchmanni* and *X. malinche* is thermal environment. To compare the lower thermal limits of fry between species, we used an ecologically-relevant quantitative measure of cold temperature tolerance for ectotherms called the critical thermal minimum, or $CT_{min}$ (Cowles & Bogert, 1944), to compare cold tolerance between newborn *X. malinche* and *X. birchmanni* fry. The $CT_{min}$ is recorded as the temperature at which a fish enters cold narcosis and is unable to move (Cowles & Bogert, 1944). To control for plasticity in thermal tolerance, 17 *X. birchmanni* and 18 *X. malinche* fry (3 broods each) were collected from mothers housed in common conditions at 22.0°C. For each species, 6-10 fry of mixed sex and aged 3-10 days were tested per $CT_{min}$ trial for a total of three trials. Two 3L plastic tanks, one per species, was filled with ambient temperature fish water (22.0°C) and placed into a secondary container. Fish water was cooled to 5°C and pumped from a cooler into each fry tank with a peristaltic pump at a flow rate of 50mL/min. Overflow was collected by the secondary container. A standard ethanol thermometer and a digital thermometer were used to monitor a standardized ramp-down rate of 0.3°C/min until the fish stopped moving entirely (following Becker & Genoway, 1979). Upon narcosis, the time and temperature were recorded, and the fry was placed in an ambient temperature tank for recovery. We fit a linear model (R stats::lm) for $CT_{min}$ with trial start temperature and date fry was born (which is the best alternative to an exact brood ID) as covariates selected with R step::step and compared means with an ANOVA and Tukey post-hoc test with R emmeans::emmeans. Statistical comparisons were See Table S19 for fry metadata and $CT_{min}$.

Though $CT_{min}$ is significantly impacted by the chosen covariates, we found that the qualitative pattern of $CT_{min}$ differences between species was the same whether we plot raw values or partial residuals of $CT_{min}$ that correct for these covariates. Therefore, we chose to plot the raw values of $CT_{min}$ of each fry so as to show the real $CT_{min}$ distribution of each species (Fig. S18).

Unexpectedly, we found that newborn *X. malinche* fry were less cold tolerant (i.e. had higher $CT_{min}$) than newborn *X. birchmanni* fry reared under common conditions (Fig. S18; p-value=0.002). However, we note that in the interest of raising fry in common conditions, fry from both species were born and reared at a temperature warmer than optimal for *X. malinche* (23°C versus 18-20°C), which may have impacted their thermal limits.

### Supplementary Figures

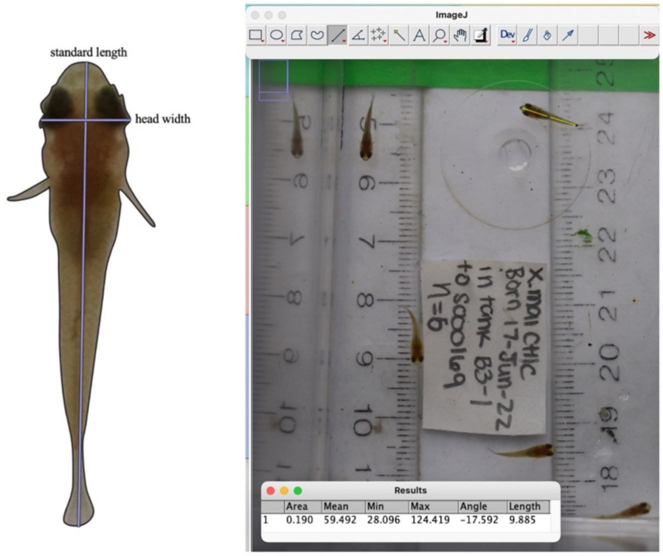


**Fig. S1.** Example of phenotyping approach for newborn fry. Tanks were checked twice daily for fry, and fry were collected immediately and photographed from above with a ruler for scale. Photos were imported to ImageJ2 and measurements were made using the line measurement tool. Standard length is a body length measure from the snout to the edge of the caudal peduncle, and head width is a measure of the widest part of the head from eye to eye.

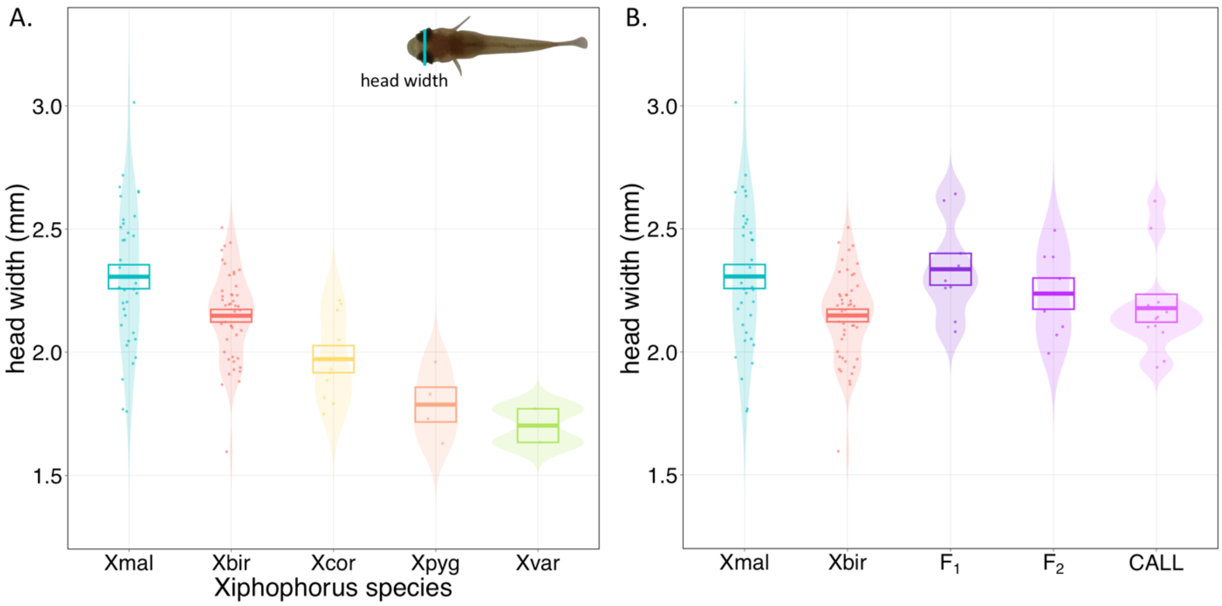


**Fig. S2.** Violin plots of head widths of newborn fry from **A.** five species: *X. birchmanni, X. malinche, X. cortezi, X. pygmaeus*, and *X. variatus* and **B.** *X. malinche*, *X. birchmanni*, $F_1$ hybrids, $F_2$ hybrids, and natural hybrids ("CALL" – Calnali Low hybrid population). All fry measured were born to lab-raised mothers in common conditions. Each data point represents the average standard length of newborn fry from one brood, colored by group. For each group, boxes show mean ± 1 standard error.

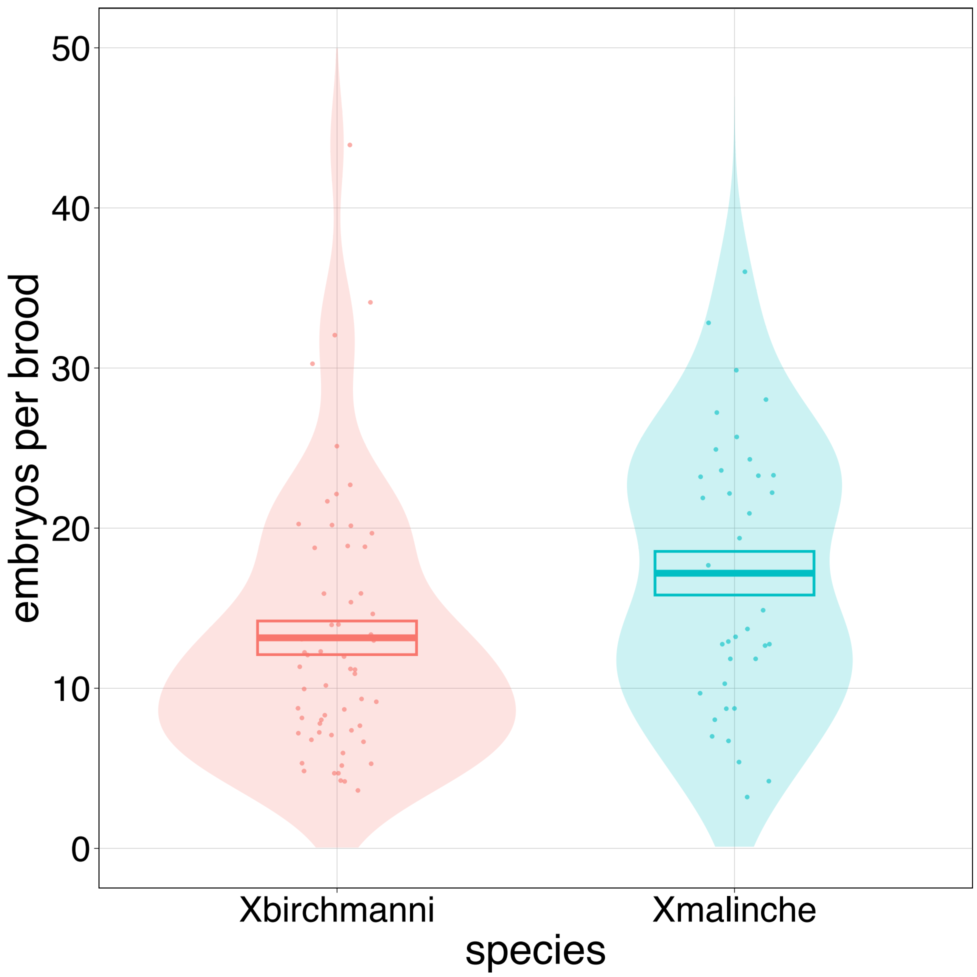


**Fig. S3.** Violin plots showing brood size measurements across wild-caught collections of *X. birchmanni* and *X. malinche* over multiple seasons. The overlayed boxes show mean brood size ±1 standard error.

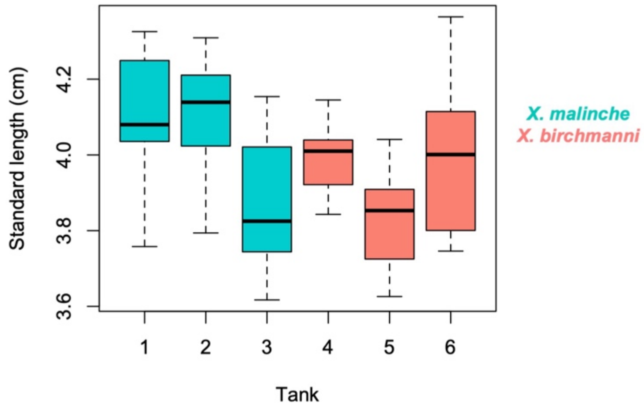


**Fig. S4.** Boxplots showing adult female size for *X. malinche* and *X. birchmanni* individuals reared in common garden conditions. On average, *X. malinche* females in this data set were slightly larger than *X. birchmanni* females, but the variation in female size between tanks exceeded the variation between species.

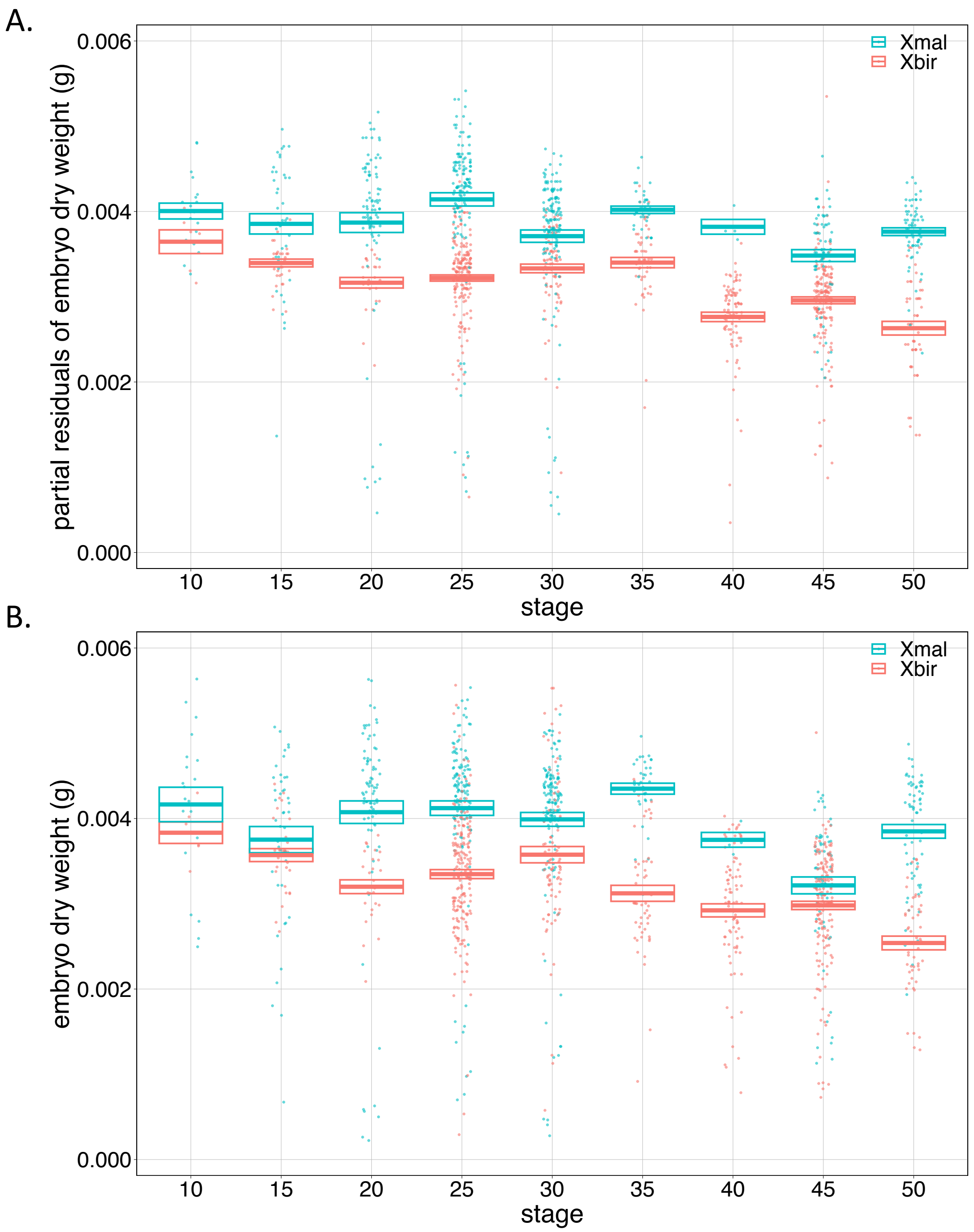


**Fig S5. A)** Developmental profiles of *X. birchmanni* (Xbir – pink) and *X. malinche* (Xmal – blue) throughout development using partial residuals of embryo dry weights. This is the same plot as in Fig. 2B of the main text but zoomed out to show all data points. Developmental stage is plotted on the x-axis and partial residuals of embryo dry weight is plotted on the y-axis. Points show the average dry weight of embryos in one brood, colored by species. For each species, boxes show mean ±1 standard error. **B)** Developmental profiles of *X. birchmanni* (Xbir – pink) and *X. malinche* (Xmal – blue) throughout development using raw embryo dry weights (rather than partial residuals of embryo dry weight; see Fig S5A and main text). Developmental stage is plotted on the x-axis and embryo dry weight is plotted on the y-axis. Points show the average dry weight of embryos in one brood, colored by species. For each species, boxes show mean ±1 standard error.

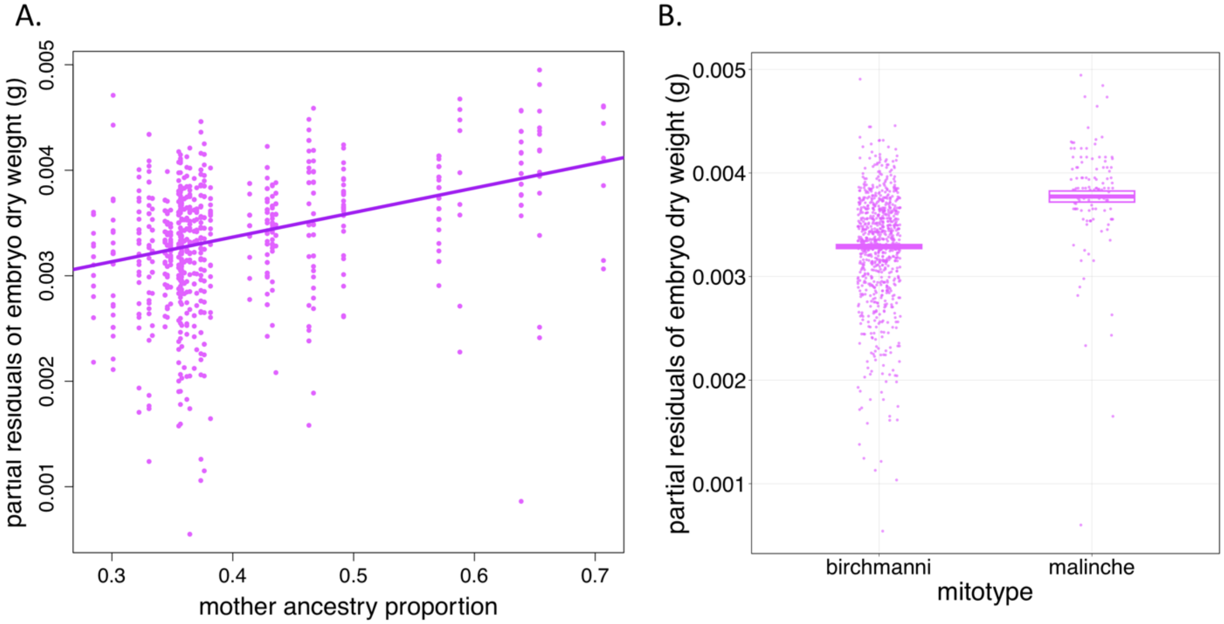


**Fig. S6.** Effects of **A.** maternal admixture proportion and **B.** maternal (and embryo) mitochondrial ancestry on embryo size (with partial residuals of embryo dry weight plotted to account for brood size as a fixed effect and brood ID as a random effect). *X. malinche* maternal admixture proportion is significantly positively correlated with embryo dry weight (L-ratio=61.6, p-value<0.0001, R=0.37), and similarly, *X. malinche* mitochondria predict larger embryos (L-ratio=11.8, p-value=0.0006). Points show the partial residuals of dry weight for each hybrid embryo. Boxes in **B** show the mean ± 1 standard error.

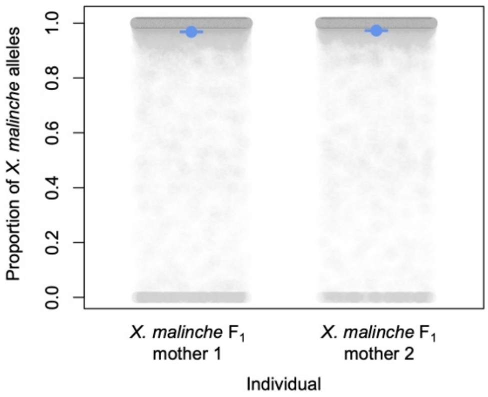


**Fig. S7.** Allele-specific expression in RNAseq data from the ovaries of *X. malinche* mothers pregnant with $F_1$ embryos. Gray points show the ratio of the *X. malinche* to *X. birchmanni* allele at ~61,000 individual ancestry informative sites across ~12,000 genes with ancestry informative sites expressed in ovarian tissue. Blue points and whiskers show the mean ratio ± 2 standard errors.

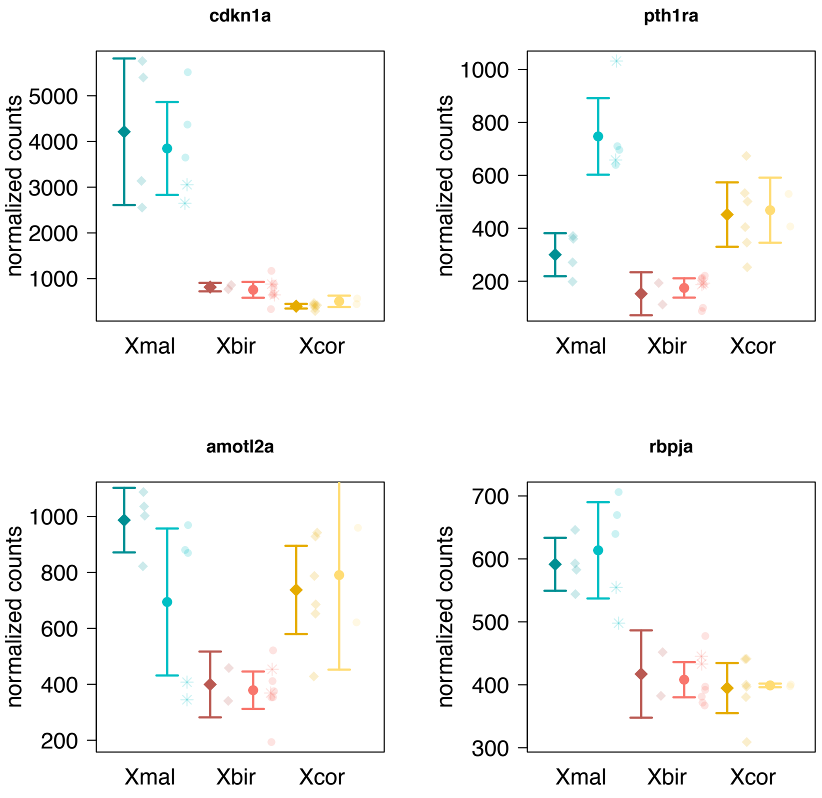


**Fig. S8.** Normalized counts generated by kallisto for a subset of genes in an artery development Gene Ontology biological pathway that was found to be significantly enriched among the genes differentially expressed between species. Species are color coded with *X. malinche* samples in blue ("Xmal"), *X. birchmanni* samples in pink ("Xbir"), and *X. cortezi* samples in yellow ("Xcor"). For each species, the two sets of counts represent the average normalized counts ± 2 standard errors for early pregnancy samples (left; in the darker color, diamond points) and late pregnancy samples (right; in the lighter color, circle and star points). Estimates from individual samples are plotted as points, except for the two wild-caught *X. malinche* and *X. birchmanni* samples, which are plotted as stars.

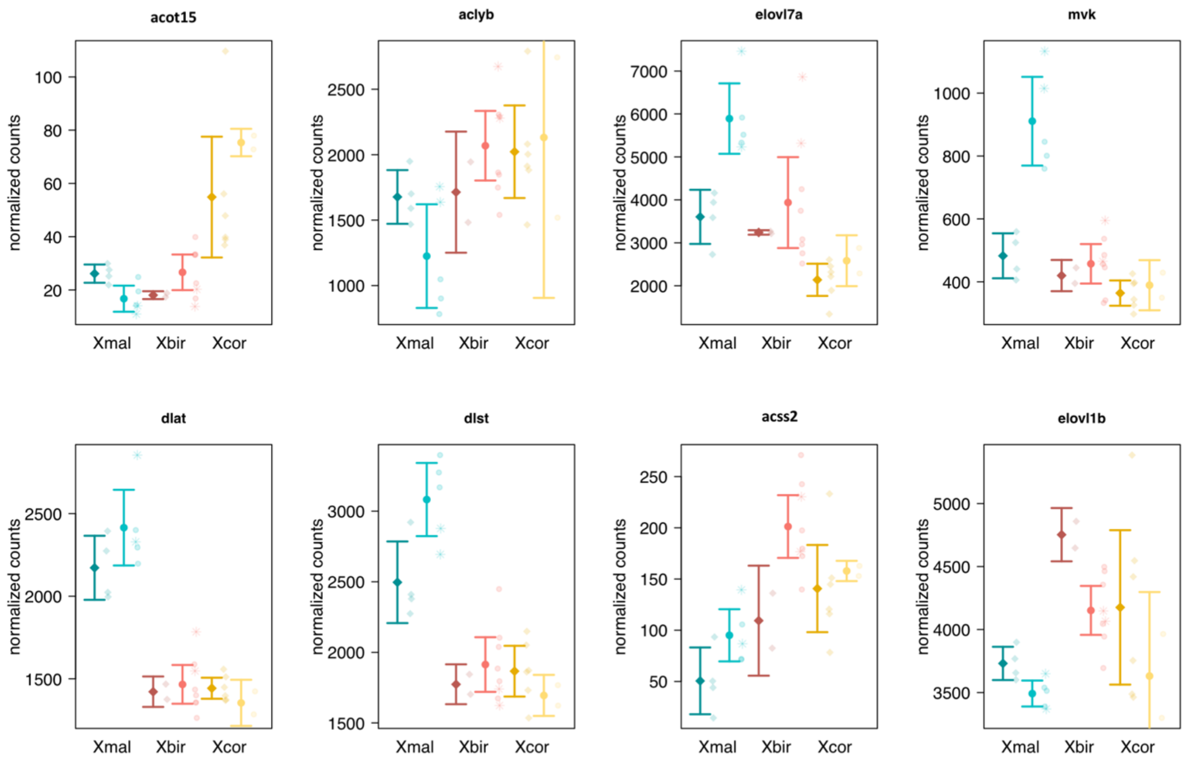


**Fig. S9.** Normalized counts generated by kallisto for a subset of genes in the acyl- and acetyl-CoA metabolic process pathways. The genes shown here were part of a WGCNA module ("darkorange2"), which strongly correlated with samples from the late pregnancy *X. malinche* ovarian expression dataset. Species are color coded with *X. malinche* samples in blue ("Xmal"), *X. bichmanni* samples in pink ("Xbir"), and *X. cortezi* samples in yellow ("Xcor"). For each species, the two sets of counts represent the average normalized counts ± 2 standard errors for early pregnancy samples (left; in the darker color, diamond points) and late pregnancy samples (right; in the lighter color, circle and star points). Estimates from individual samples are plotted as points, except for the two wild-caught *X. malinche* and *X. birchmanni* samples, which are plotted as stars.

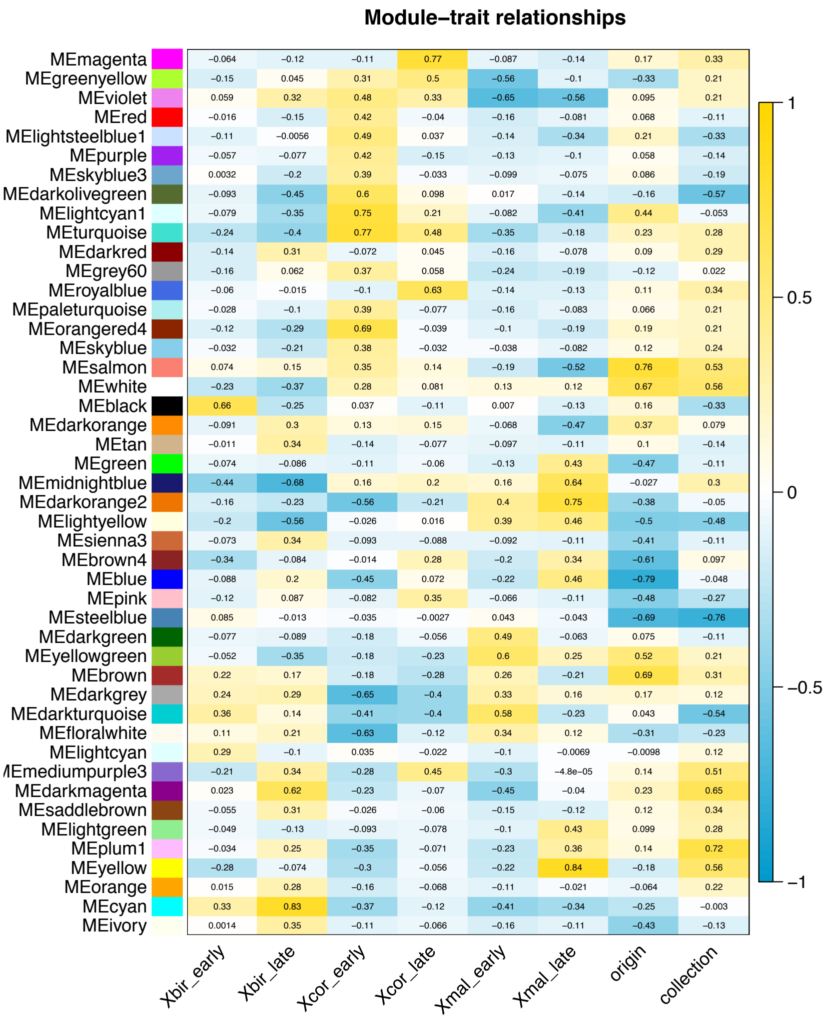


**Fig. S10.** Heat map showing module-trait relationships for co-expression modules identified by WGCNA analysis of ovarian RNAseq data (see main text). The arbitrary module name is plotted on the y-axis and the biological or technical variable being analyzed is plotted on the x-axis. Each square shows the Pearson’s correlation between the expression eigenvector for that module and the variable of interest (e.g species and developmental stage). The color bar from blue to yellow shows the value and direction of the Pearson’s correlation. For example, module “MEmagenta” is strongly positively correlated with late-stage *X. cortezi* embryos (R=0.77).

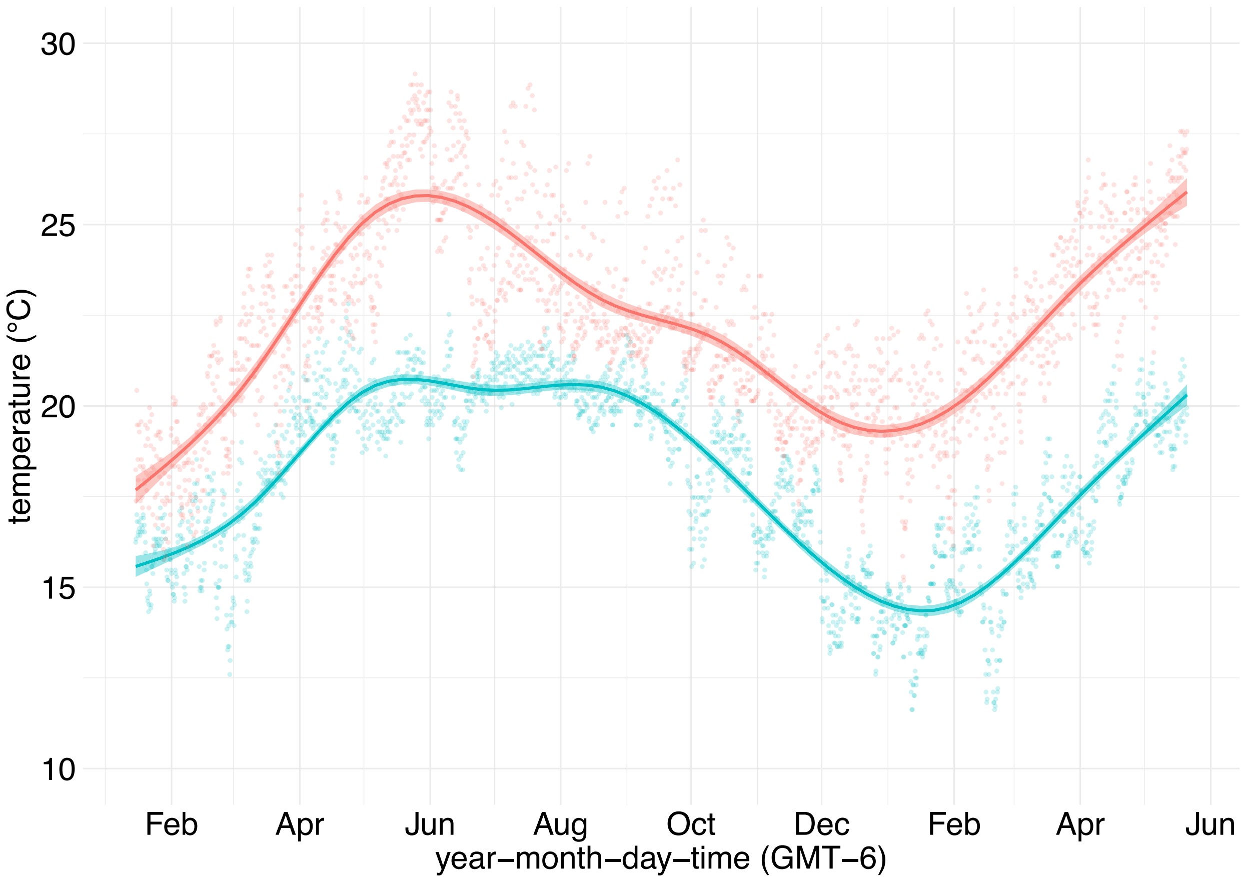


**Fig. S11.** Temperature differences between *X. birchmanni* and *X. malinche* typical elevations over one year. Data is reproduced from (Payne et al., n.d.), showing temperature data collected by HOBO loggers at the *X. malinche* Chicayotla population from February 2020 – June 2021 (in blue), and at the Acuapa population, an introgressed population found at *X. birchmanni*-typical elevations, from February 2016 – February 2017 (in pink).

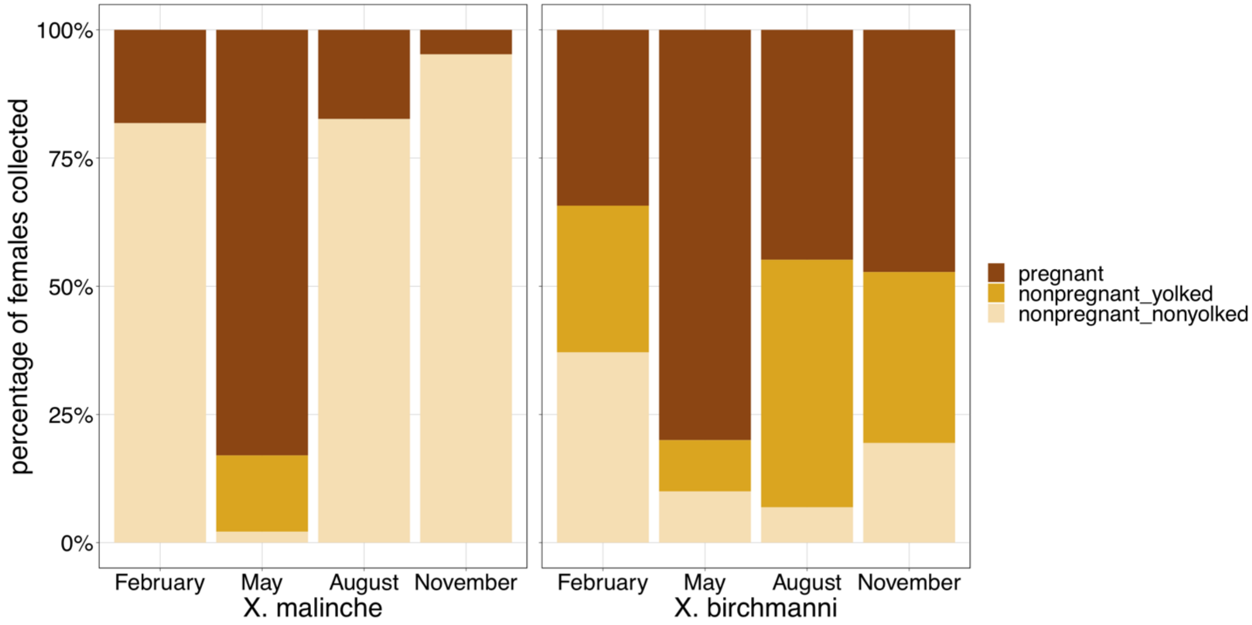


**Fig. S12.** Proportion of collected females that fell into different reproductive categories across seasonal collections from *X. malinche* (left) and *X. birchmanni* (right) populations from 2020-2023. While *X. malinche* females exhibit strong seasonality in whether they had mature (yolked) eggs, or were pregnant, we saw less seasonality in *X. birchmanni*, with pregnant females observed across all seasons.

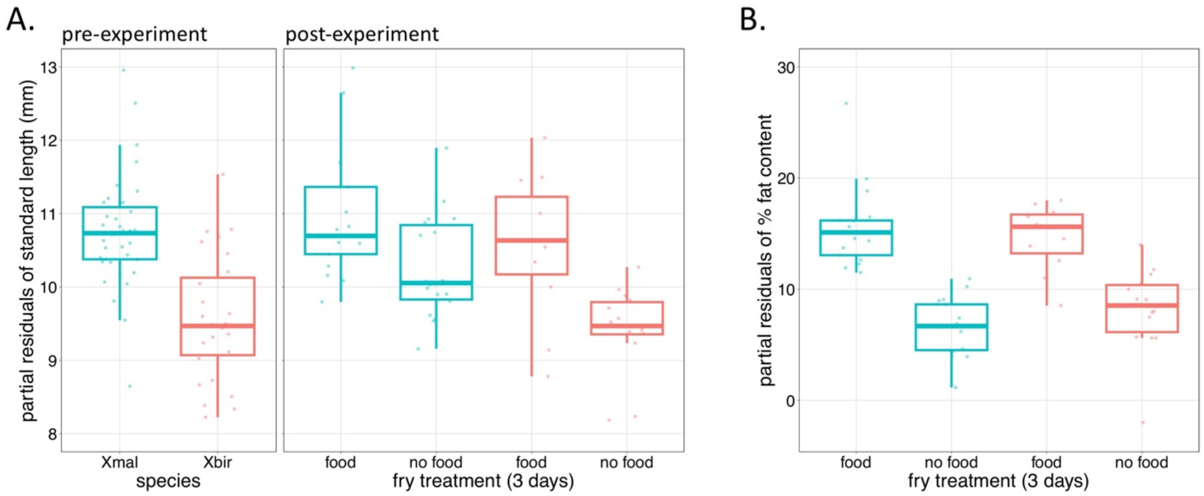


**Fig. S13.** Boxplots of partial residuals of **A**) standard length and **B**) % fat content for newborn *X. malinche* and *X. birchmanni* fry that experienced either food or no food treatments for three days. (**A**)While both species lost length in the no food treatments, *X. birchmanni* (pink) lost more length than *X. malinche* (pink). By contrast, while fry from both species gained less fat when starved, there was no significant difference in fat content between species in either condition (**B**). However, we found that starved *X. birchmanni* had significantly lower body weight than *X. malinche* (Fig. 4).

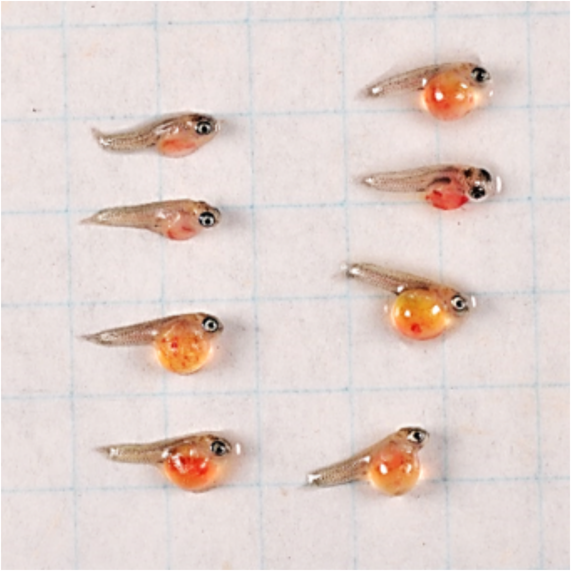

**Fig. S14.** Example of premature birth phenotype for one brood born to an *X. birchmanni* mother crossed to an *X. malinche* father. Note the unabsorbed egg sac in all individuals. This is the typical developmental stage at which we observe premature birth in this cross direction (Table S24).

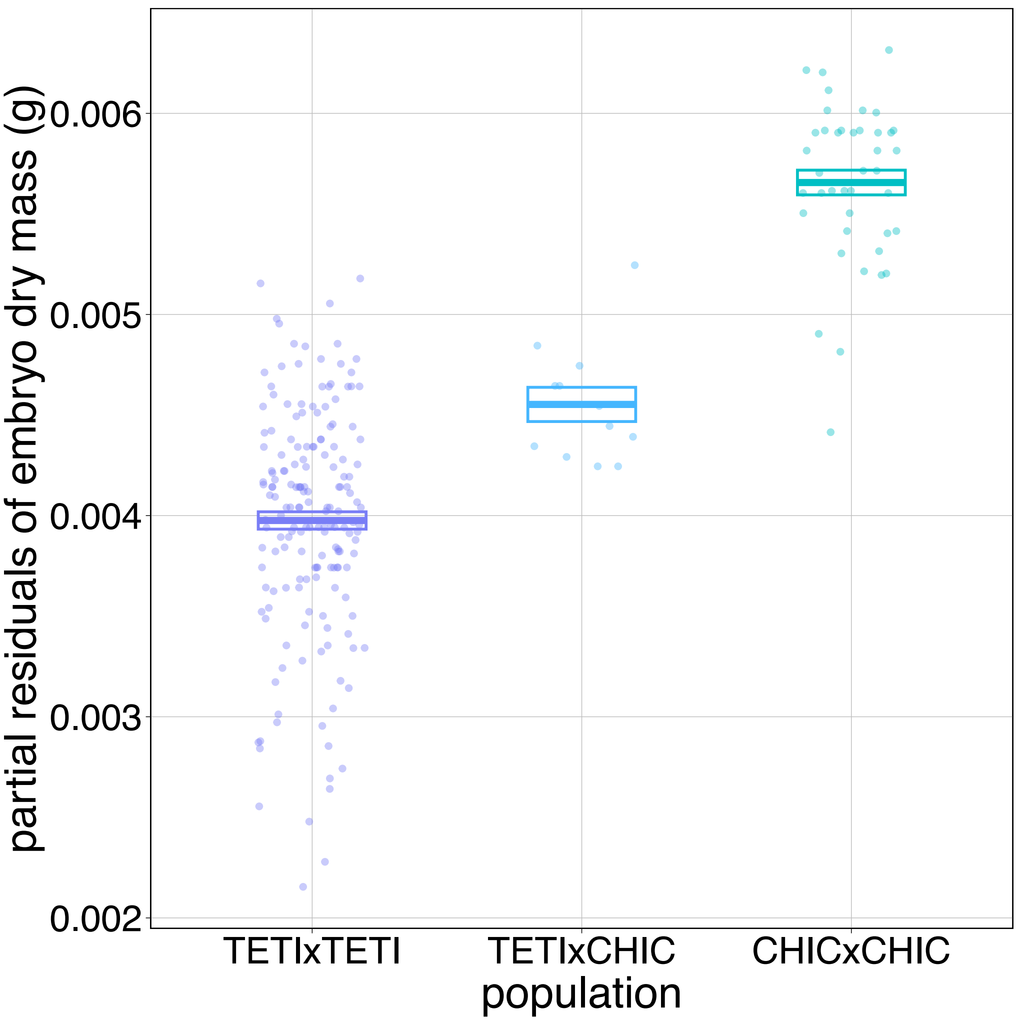


**Fig. S15.** Boxplots of partial residuals of embryo dry weights from within and between population crosses of the allopatric *X. malinche* populations Chicayotla and Tetipanchalco. Crosses were performed with artificial insemination in lab, dissected and developmentally staged, before being dried and weighed as described in the main text. For this analysis, only crosses with embryos between stages 25 and 35 are compared as the TETIxCHIC cross only yielded stage 25 and 35 embryos.

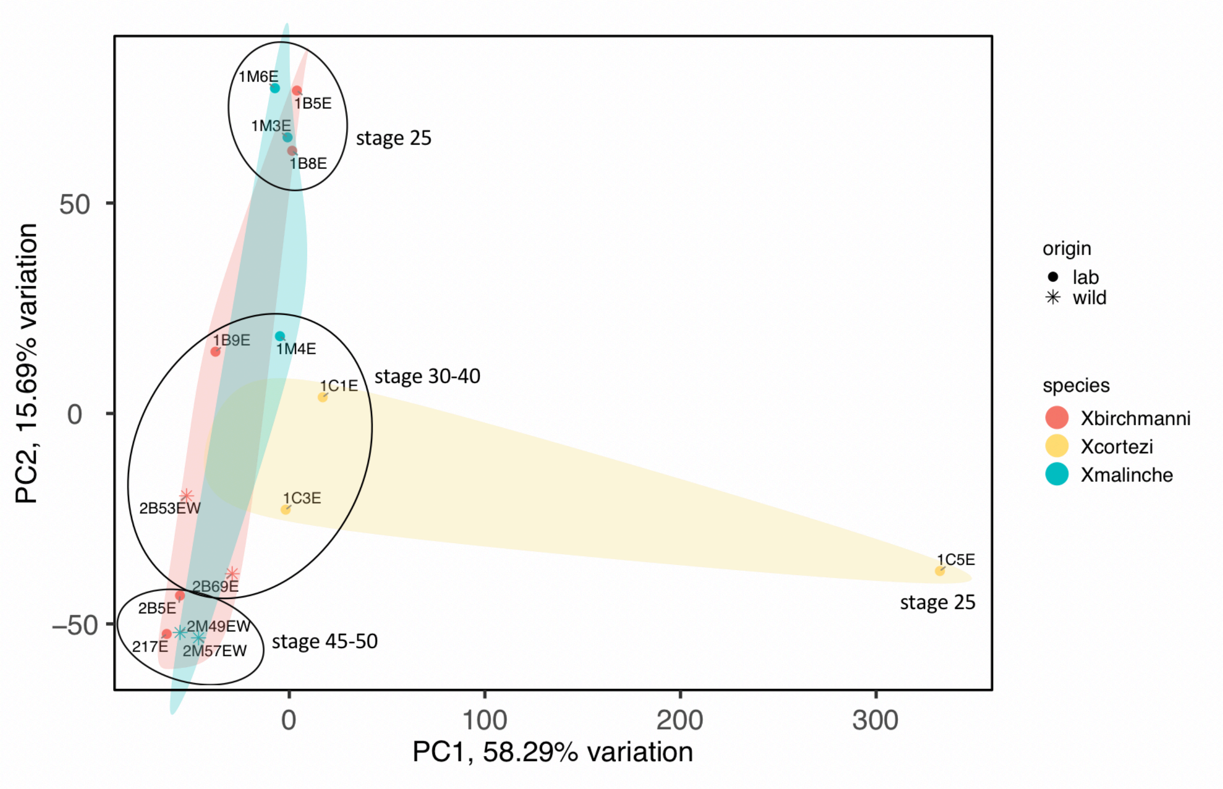


**Fig. S16.** Principal component analysis of transformed gene abundance counts from RNAseq data of mid- to late-stage embryos from *X. birchmanni* (pink), *X. malinche* (blue), and *X. cortezi* (yellow). For each of *X. birchmanni* and *X. malinche,* two wild-caught samples were also included in the dataset (denoted by asterisks). Samples cluster by developmental stage, and *X. cortezi* separates from *X. birchmanni* and *X. malinche* on PC1. Black circles label the developmental stages represented in each cluster. Colored envelops show the space in PC1 and PC2 occupied by all samples of a given species.

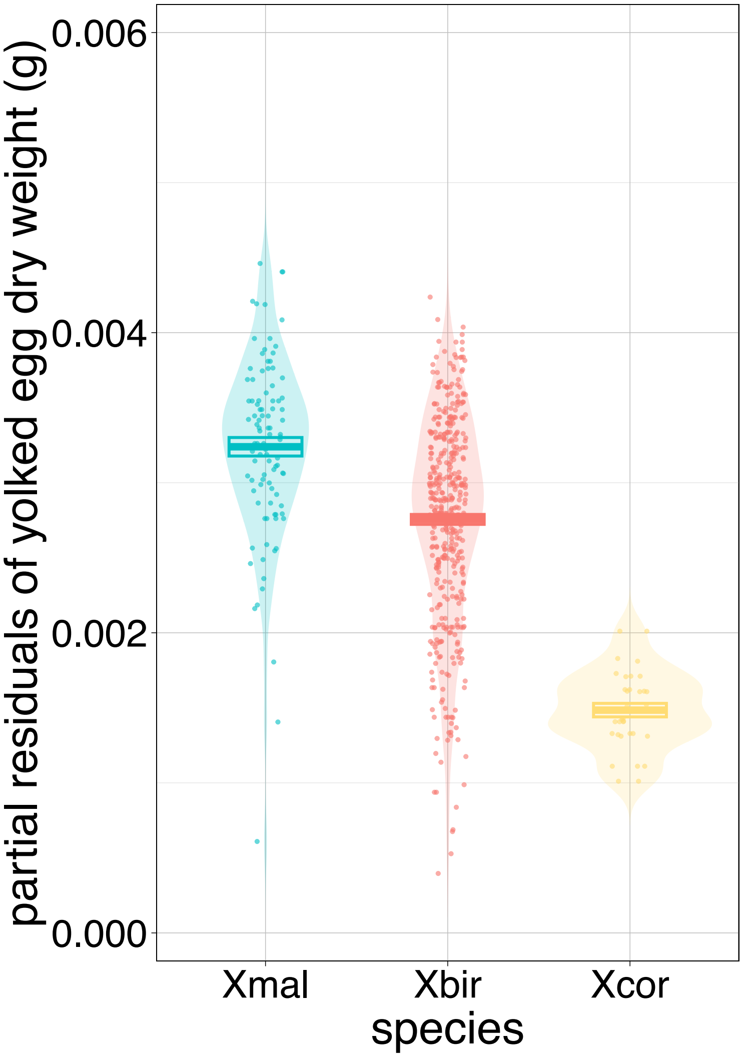


**Fig. S17.** Violin plot comparing stage zero (fully-yolked but unfertilized) eggs between *X. birchmanni, X. malinche*, and *X. cortezi*. Points show the partial residuals of average egg dry weight per brood and boxes show mean and ± 1 standard error. These profiles were generated from wild-caught fish.

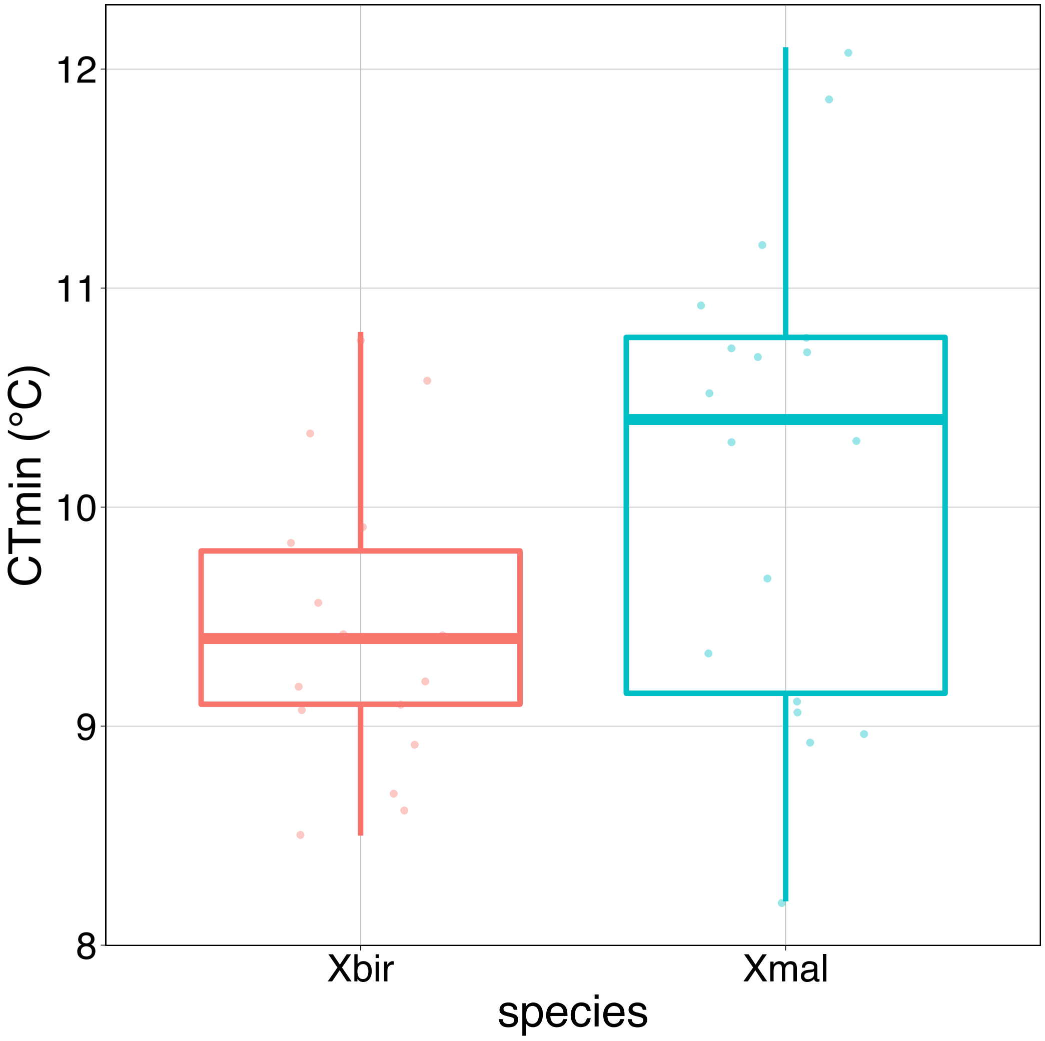


**Fig. S18.** Boxplots of critical thermal minimum ($CT_{min}$) for newborn *X. malinche* (blue) and *X. birchmanni* (pink) fry. Unexpectedly, newborn *X. malinche* fry were less cold tolerant (i.e. had higher $CT_{min}$) than newborn *X. birchmanni* fry reared under common conditions (p-value=0.002). However, we note that in the interest of raising fry in common conditions, fry from both species were born and reared at a temperature warmer than optimal for *X. malinche* (23°C versus 18-20°C), which may have impacted their thermal limits (see Supplementary Information 11).
